# Rhodopsin 7 is indispensable for regulating the firing rates of olfactory sensory neurons in response to extracellular field potential changes in *Drosophila melanogaster*

**DOI:** 10.64898/2025.12.05.692481

**Authors:** Masaki Kataoka, Keisuke Saito, Kazuaki Ikeda, Hiroshi Ishikita, Nobuaki K. Tanaka

## Abstract

Although extracellular field potential changes are commonly observed in the nervous systems, it remains controversial if extracellular electrical activity contributes to neural processing or whether it is an epiphenomenon associated with neural activity. We previously reported that the extracellular field potential change in compound eyes in response to light stimulation induces firing rate changes in olfactory sensory neurons in female *Drosophila melanogaster*. Through further investigation, we found that the extracellular field potential within the olfactory sensillum is regulated by octopaminergic neurons in response to the light stimulation and that rhodopsin 7 mediates the firing rate changes in the olfactory sensory neurons in response to field potential changes in a light-independent manner. Structural analysis suggests a voltage-dependent gating mechanism for rhodopsin 7 to respond to the field potential change. This study reveals that the nervous system actively controls the field potential in response to sensory input, resulting in alteration of behavioral patterns as well as neural firing patterns in a context-dependent manner.

**Significance statement:** Although extracellular electrical activity has been recorded to diagnose neuropsychiatric disorders, it remains uncertain how it can be controlled by the nervous system. Moreover, it is difficult to investigate how neurons change their excitability by responding to the change in the extracellular field potential, as synaptic communication interferes in the ability to isolate the function of extracellular electrical activity. We here show that the extracellular field potential within the olfactory sensillum in *Drosophila melanogaster* is actively regulated by octopaminergic neurons in response to sensory input. We also provide evidence that rhodopsin, a major light sensor protein, mediates responses to extracellular electrical signals, resulting in alternation of behavioral patterns as well as neural firing patterns in a context-dependent manner.

## Introduction

The extracellular field potential changes continuously in many nervous systems (Buzsáki et al., 2012; Weiss and Faber, 2010). The most representative of these field potential changes are those observed in electroencephalograms in mammals and odor-evoked neural oscillations in insects (Stopfer et al., 1997; Tanaka et al., 2009; Buzsáki et al., 2012). Although the extracellular field potential has generally been considered a side effect of neuronal spiking, recent studies have demonstrated that the electric field effects generated by the changes in extracellular field potential can affect neural processing. Ephaptic transmission is a form of communication between neighboring neurons that does not depend on synapses (Anastassiou and Koch, 2015). For example, excitation of one olfactory sensory neuron (OSN) induces ephaptic inhibition of a nearby OSN housed in the same sensillum in *Drosophila* antennae (Su et al., 2012; Zhang et al., 2019), and firing of a Purkinje cell promotes synchronous firing of nearby Purkinje cells by ephaptic coupling (Han et al., 2018). Furthermore, externally imposed electric fields created by injecting current into a cortical slice affect the synchronicity of neural firings (Anastassiou et al., 2011). Although these studies have indicated that extracellular electrical activity possesses an important role in neural processing, it remains uncertain whether it can be controlled by the nervous system. Moreover, it is difficult to investigate how neurons change their excitability by responding to the change in the extracellular field potential in the naive nervous system, as synaptic communication between neurons is always present in vivo and interferes in the ability to isolate the function of extracellular electrical activity.

We previously reported that light-evoked firing rate increases in the OSNs of *Drosophila melanogaster* are mediated primarily by ephaptic transmission of light information from the retinal photoreceptor cells (Ikeda et al., 2022). To reveal the ephaptic transmission, we recorded the odor responses from the trichodia (T1) sensilla (Fig. 1A). The T1 sensillum is a good model to reveal the functional role of extracellular electrical activity because it houses a single olfactory sensory neuron, and the extracellular fluid within the sensillum is segregated from the hemolymph by septate junctions (Shanbhag et al., 2000). These features enabled us to investigate the role of the extracellular field potential in neuronal excitability by manipulating it without inducing synaptic communication between neurons within the sensillum. In this study, we further investigated how the ephaptic transmission of light information from the retinae results in increased firing rates in the OSNs of the distant T1 sensilla. We identified photoreceptor proteins and efferent neurons indispensable for this sensory integration, revealing that the nervous system has an active mechanism to control the firing rates of neurons by changing the extracellular field potential in response to sensory input.

**Figure 1.**
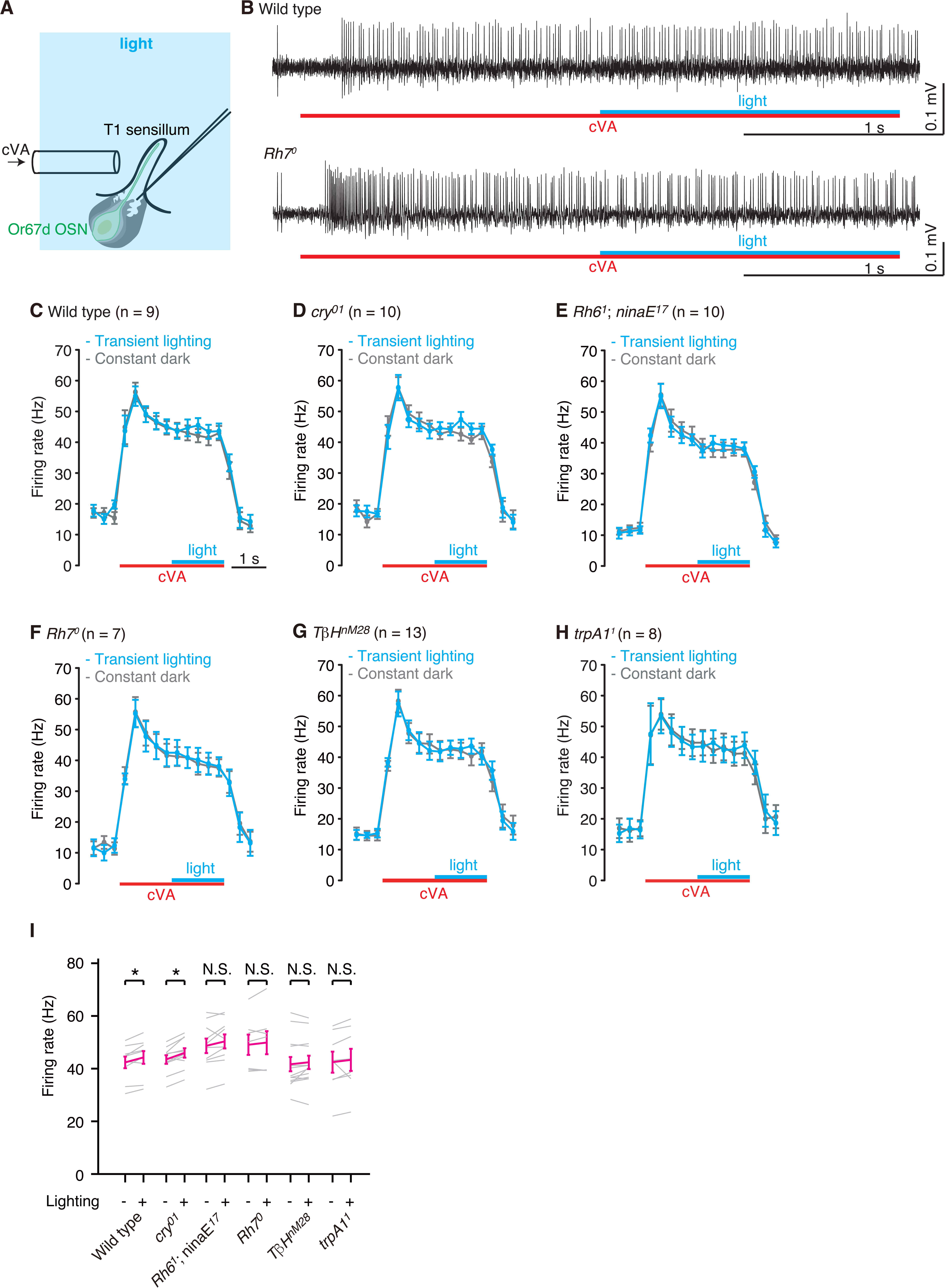
Transient light stimulation during the latter half of *cis*-vaccenyl acetate (cVA) response increased the firing rates of the Or67d-expressing olfactory sensory neurons (OSNs). ***A***, Experimental setup. While recording spikes extracellularly from Or67d-expressing OSN in a T1 sensillum, *cis*-vaccenyl acetate (cVA) and blue light stimulations were applied to the fly. The T1 sensillum is composed of the Or67d-expressing OSN (green) and auxiliary cells (gray). ***B*,** The Or67d-expressing OSN in a T1 sensillum responds specifically to cVA in *Rh7^0^* mutants as well as wild type. ***C*-*H*,** A transient light pulse applied during the latter half of cVA stimulation significantly increased the firing rates of Or67d-expressing OSNs in wild type (***C***) and *cry^01^*(***D***), but not in *Rh6^1^*; *ninaE^17^*(***E***), *Rh7^0^* (***F***), *TβH^nM28^* (***G***), and *trpA1^1^* (***H***) mutant flies. Each panel shows the firing rates in each bin of 300 ms. Cyan plots represent the firing rates of the trials in which both odor and transient light stimuli were applied, whereas gray plots show those where only odor stimuli were applied in darkness. Horizontal cyan lines represent 1.5 s of light stimulation, whereas red lines indicate the 3-s odor stimulation. ***I*,** The firing rates during the latter half of the cVA stimulation. Each gray line represents the firing rate of a different animal, and magenta represents the mean. Error bars indicate the standard errors of the mean. For statistical analyses, Wilcoxon matched-pairs signed rank tests were performed. *, *p* < 0.05; N.S., not statistically significant. See Table 1 for full statistical reporting.

## Materials and methods

### *Drosophila* strains

Flies were reared on standard agar-cornmeal medium at 25°C with a 12-hour light/12-hour dark cycles unless noted. We used female flies of Canton S as a wild type and the following mutants 3 to 7 days after eclosion: *eyes absent* (*eya^2^*) (RRID:BDSC_2285; Choi and Benzer, 1994), *Rh7^0^*(RRID:BDSC_83716; Kistenpfenning et al., 2017), *no receptor potential A^36^* (*norpA^36^*) (RRID: BDSC_9048; Bloomquist et al., 1988), and *transient receptor potential channel A1^1^* (*trpA1^1^*) (RRID: BDSC_36342; Kwon et al., 2008) obtained from the Bloomington *Drosophila* Stock Center (BDSC); *Rh6^1^*; *ninaE^17^* (Hanai et al., 2008) from Kyoto *Drosophila* Stock Center; *cry^01^* (Dolezelova et al., 2007) gifted from Fumika Hamada; and *TβH^nM28^* (Certel et al., 2007) from Sarah Certel. We prepared flies in which synaptic transmission of the photoreceptors in the ocelli, compound eyes, and eyelets were blocked by mating *longGMR-GAL4* (RRID:BDSC_8121; Wernet et al., 2003) from BDSC with *UAS-tetanus toxin* (TNT) (RRID:BDSC_28838; Sweeney et al., 1995) from Aki Ejima. Cell death and down-regulation of rhodopsin 7 (Rh7) expression were induced by *UAS-DTI* (RRID:BDSC_25039; Bellen et al., 1992; Han et al., 2000) and *UAS-Rh7-RNAi* (RRID:BDSC_62176; Perkins et al., 2015) from BDSC, respectively. These two strains were mated with *Or67d-GAL4* line (Fishilevich and Vosshall, 2005) gifted by Leslie Vosshall to express the transgene specifically in the Or67d-expressing OSNs. Cell death by *DTI* was induced by maintaining newly eclosed flies at 29°C for five days (Berdnik et al., 2006) and then at 25°C for one or two days before performing the experiments.

Flies bearing both *UAS-GFP* (obtained from Barry Dickson; Tanaka et al., 2008) and *UAS-20XmCD8::GFP* (from BDSC; RRID:BDSC_32194) were crossed with the following GAL4 strains to label aminergic fibers: *NP7088 GAL4* enhancer-trap strain (from Kyoto *Drosophila* Stock Center; Tanaka et al., 2008) and *Tdc2-GAL4* line (from BDSC; RRID:BDSC_9313; Cole et al., 2005; Busch et al., 2009) for labeling octopaminergic neurons; *Hn.493-* and *Hn.819-GAL4* strains (from BDSC; RRID:BDSC_66009 and 66010; Lee et al., 2011) for serotonergic neurons; *Ddc-*(*HL7-71* and *HL9-61*) and *TH-GAL4* lines (from Taro Ueno and Kazuhiko Kume; Claridge-Chang et al., 2009; Ueno et al., 2012) for dopaminergic neurons.

All animal experiments were performed in accordance with the relevant guidelines and regulations of Hokkaido University.

### Electrophysiology

A female fly was anesthetized in a vial on ice for less than 1 min and restrained in a custom-made plastic dish by fixing the appendages with wax and epoxy (Tanaka et al., 2009). The right antenna was then set on a tin foil to which the rear side of the antenna was bonded with epoxy to avoid movement. After a small window was opened in the thorax by partially removing the cuticle, *Drosophila* saline (in mM: NaCl 103, KCl 3, N-tris(hydroxymethyl)methyl-2-aminoethanesulfonic acid 5, trehalose 10, glucose 10, sucrose 7, NaHCO_3_ 26, NaH_2_PO_4_ 1, CaCl_2_ 1.5, MgCl_2_ 4, adjusted to pH 7.25 with HCl) was applied over the thorax (Ikeda et al., 2022). For the octopamine application, 5 μl of saline or 1 mM octopamine in saline was applied to 20 μl of saline covering the window of the thorax.

The restrained flies were then placed under a Slicescope (Scientifica, East Sussex, UK) equipped with a GSWH20x ocular lens, LUCPlanFLN40x objective, and U-ECA 2x magnification changers (Olympus, Tokyo, Japan). The recording and reference electrodes were inserted into the sensillum through the socket (T1 sensillum) or bristle cuticle (ab2) and saline over the thorax, respectively, except for the octopamine application experiments for which a reference electrode was inserted into the ventromedial side of the antenna to directly measure the transepithelial potential or placed on the compound eye.

We recorded neural responses from a single sensillum or a compound eye in each animal. For the sensillum recording, the recording electrodes (impedance of approximately 90 MΩ) were prepared by pulling quartz capillaries (QF100-70-7.5, Sutter Instrument, Novato, CA, USA) with a Sutter Instrument P2000 puller and were filled with sensillum lymph ringer (in mM: KCl 171.9, KH_2_PO_4_ 9.2, K_2_HPO_4_ 10.8, MgCl 3, CaCl_2_ 1, glucose 22.5, NaCl 25, adjusted to pH 6.5 with HCl) (Kaissling and Thorson, 1980; Dobritsa et al., 2003; Olsson and Hansson, 2013). Reference electrodes and recording electrodes for electroretinogram (approximately 35 MΩ) were made of glass capillaries (BF100-58-10, Sutter Instrument) with a Sutter Instruments P1000 puller and were filled with the *Drosophila* saline. To induce 0.97 - 2.2-mV LFP deflections, we injected a 5- or 8-pA current into the sensillum lymph for 1 s during the odor stimulations.

Recording signals were amplified by 100 folds with a headstage (8024/7001 N=1, Dagan Corporation, Minneapolis, MN, USA) and a DAGAN 8700 Cell Explorer amplifier or by 10 folds with a CV-7B headstage and a Multiclamp 700 B (Molecular Devices, San Jose, CA, USA) and were fed into a PC via an A/D converter Digidata 1322A or 1440A (Molecular Devices). Data were acquired digitally at a sampling rate of 5 kHz using Axoscope 10.7 or Clampex 10.7 software (Molecular Devices).

All recorded data were analyzed using MATLAB R2021b (The MathWorks, Inc., Natick, MA, USA). For the single-sensillum and LFP recordings, we high-pass filtered the recorded signal at 65 Hz and low-pass filtered at 30 Hz, respectively. We plotted all the recorded data to confirm that all the spikes were counted correctly.

### Odor and light stimulations

For the single odor stimulations, an air puff (3 s, 240 mL/min) from a pneumatic picopump (PV 820, World Precision Instruments, Sarasota, FL, USA) that passed through a tube inserted with filter paper (3.5 x 50 mm) containing 15 μL of either paraffin oil or 1% v/v cVA in paraffin oil was applied to a fly every 30 s except for the current injection and octopamine application experiments where the odorized air was puffed every 15 s. The airpuff speed was adjusted to 1 m/s. For the odor mixture experiments, an air puff (1.5 s, 500 mL/min) that passed through a vial containing paraffin oil or hexanol diluted in paraffin oil was injected into an air flow (3 s, 500 mL/min) that passed through a flask containing paraffin oil or ethyl acetate diluted in paraffin oil. Odorized air was continuously removed using a vacuum tube set near the fly’s body.

Flies were also subjected to 470-495 nm light stimulations. The light was originally emitted from an Olympus U-HGLGPS mercury lamp and then was filtered by U-MNIBA3. The light intensity was adjusted to 55 μmol m^-2^ s^-1^ except for the case of *trpA1^1^* (37 μmol m^-2^ s^-1^), which was confirmed by a LI-250 light meter connected to an LI-190SA quantum sensor (LI-COR, Lincoln, NE, USA). We alternated the order of different light or odor conditions and counterbalanced the number of recordings in which one light or odor condition preceded the other with that in the opposite order. For each condition, we measured sensory responses three or five times from each animal and averaged them for further analyses.

### Antibody staining of brains and antennae

The antennae were stained with antibodies as follows: First, flies were vortexed in 100% ethanol and washed with phosphate buffered saline (PBS). The distal tips of the antennae were cut off to create a hole. The antennae were then amputated, fixed in 4% paraformaldehyde (PFA) for 1 h, washed in PBS, and shaken in 0.2% triton in PBS (PBST) for 2 h. They were incubated in blocking solution (10% goat serum in 1% PBST) containing primary antibodies for two days at room temperature with shaking. After washing with 0.2% PBST, they were incubated in blocking solution containing secondary antibodies for one day. After washing with PBS, they were added to 50% glycerol in PBS and then mounted in Vectashield mounting medium (Vector Laboratories, Inc., Burlingame, CA, USA).

Brains were dissected out in PBS and fixed in 4% PFA in PBS for 40-60 min. After washing with PBS, the brains were shaken in the blocking solution. They were incubated in the blocking solution containing primary antibodies overnight at room temperature with shaking. After washing with 0.2% PBST, the brains were incubated in the blocking solution containing secondary antibodies overnight at room temperature with shaking. Finally, the brains were washed with PBS and mounted in 80% glycerol in PBS.

The primary antibodies used were rabbit anti-GFP (A11122, Molecular Probes, Eugene, OR; RRID:AB_221569; diluted at 1:500 or #598, Medical & Biological Laboratories, Tokyo, Japan; RRID: AB_591819; diluted at 1:500), rabbit anti-serotonin (S5545, Sigma-Aldrich, St. Louis, MO, USA; 1:125), rabbit anti-octopamine (AB-T070; Advanced Targeting Systems, Carlsbad, CA, USA; 1:200), rabbit anti-tyrosine hydroxylase (NB300-109, Novus Biologicals, Cetennial, CO, USA; 1:200) (Alekseyenko et al., 2010), rabbit anti-histamine antibodies (#22939, ImmunoStar, Hudson, WI, USA; 1:200), and rabbit anti-Rh7 antibodies (1:100) gifted from Charlotte Helfrich-Förster and Craig Montel (Kistenpfenning et al., 2017; Ni et al., 2017). Goat anti-rabbit antibodies conjugated with Alexa Fluor 488 or 647 (A11034 and A21245, Molecular Probes; RRID:AB_2576217 and 2535813; 1:500) or donkey anti-rabbit antibodies conjugated with Alexa Fluor 594 (A21207; RRID:AB_141637; 1:500) were used as secondary antibodies.

Confocal images of the immunostained brains and antennae were taken with a laser scanning confocal microscope (LSM 700, Zeiss, Jena, Germany).

### Structural analysis and molecular dynamics simulations of Rh7

Based on the amino-acid sequences of Rh1 (NP_524407.1), Rh7 (AAF49949.2), and the G-protein α subunit q (CG17759) from *Drosophila melanogaster*, five protein structures were predicted by using AlphaFold2 (Yang et al., 2023). Among the structures, the top ranked structure (rank 1) was selected for analyses. For Rh7, to investigate the influence on transmembrane potential changes, two protonation patterns were evaluated. In the standard protonation state, all titratable groups (i.e., acidic and basic groups) were ionized. Upon protonation of these acidic residues in helix 10, Glu and Asp in the intracellular region (i.e., Glu434, 443, 456, 457, 459, 460, 462, 463, and Asp439) were fully protonated.

The Rh7 assembly was embedded in a lipid bilayer consisting of 257 1-palmitoyl-2-oleyl-sn-glycero-3-phosphocholine (POPC) molecules using CHARMM-GUI (Jo et al., 2008) and soaked in 58507 TIP3P water models, and 5 and 16 chloride ions (the deprotonated- and protonated-acid patterns, respectively) were added to neutralize the system using VMD plugins (Humphrey et al., 1996). After structural optimization with position restraints on heavy atoms of the Rh7 assembly, the system was equilibrated at 300 K for 2 ns with a time step of 0.5 fs, and annealed from 300 to 0 K over 115 ps with a time step of 0.1 fs. The positional restraints on heavy atoms of side-chains were released, and the system was heated from 0.1 to 300 K over 275 ps with a time step of 0.05 fs, equilibrated at 300 K for 1 ns with a time step of 0.5 fs, and annealed from 300 to 0 K over 115 ps with a time step of 0.1 fs. The positional restraints were released except for atoms of backbones in the transmembrane region, which were treated as harmonic constraints with force constants set to 5 kcal/mol. The system was heated from 0.1 K to 300 K over 5.5 ps with a time step of 0.05 fs and equilibrated at 300 K for 1 ns with a time step of 0.5 fs. The system was equilibrated at 300 K for 108 ns with a time step of 1.5 fs. All molecular dynamics (MD) simulations were conducted with the CHARMM22 force field parameter set (MacKerell et al., 1988) using the MD engine NAMD version 2.13 (Phillips et al., 2005). For MD simulations with a time step of 1.5 fs, the SHAKE algorithm for hydrogen constraints was employed (Ryckaert et al., 1977). For temperature and pressure control, the Langevin thermostat and piston were used (Feller et al., 1995).

The Rh7/G-protein complex structure was generated by superimposing the Rh7 and G-protein structures onto the reported cryo-EM structure of the rhodopsin-transducin complex (PDB code 6OY9) (Gao et al., 2019) using the align command in PyMOL Molecular Graphics System, Version 1.2r3pre (Schrödinger, LLC.) (Supporting coordinates). Supporting coordinates of model structures were generated using AlphaFold2 and subsequently modified for figure presentation.

### Behavioral assays

Behavioral experiments were performed in dim red light at a zeitgeber time of 1-4 h at 25°C and more than 40% humidity. A double-layered courtship chamber (Ejima et al., 2007) was used to set cVA in the lower cell and a virgin female in the upper cell as follows. First, filter paper (φ = 2 mm, 0.18-mm thickness) was set on double-sided tape (0.5-mm thickness) at the center of the lower cell (φ = 8 mm, 3-mm depth, without a top) and soaked with 0.2 μL of hexane or cVA solution (18-ng cVA in hexane). After 30 min to evaporate hexane, the cell was covered with a cover glass, over which a nylon mesh and the upper cell (φ = 8 mm, 3-mm depth) were placed. A virgin female was introduced into the upper cell and allowed to acclimate to the cell for 5 min. The cover glass inserted between the lower cell and mesh was then removed to expose the female to the odor, and the resulting behavior was recorded at 15 Hz with a GZ4304M video camera (Gazo, Niigata, Japan). One minute later, a blue light stimulation was applied for 1.5 s for “+light” animals, but not for naïve “-light” animals. The centroid of the female’s body in three frames just after the light offset time was tracked with MTrack2 plugin in ImageJ Fiji (https://valelab4.ucsf.edu/~nstuurman/IJplugins/MTrack2.html) and cos(*θ*) was calculated using MATLAB, where *θ* was the angle between the unit vector joining the centroid in the first frame to the center of the cell and the one joining the centroid in the first frame to that in the third. The positive cos(*θ*) indicates that the female approaches the center of the cell, whereas the negative one shows the female leaves the center.

### Experimental design and statistical analysis

Data are presented as means ± standard errors of the mean. We performed Wilcoxon matched-pairs signed rank test, Mann-Whitney test, Wilcoxon rank sum exact test adjusted by Bonferroni correction, and Kruskal-Wallis test followed by Dunn’s multiple comparison test using Prism 10 (GraphPad, San Diego, CA, USA), R 3.3.3 (https://www.R-project.org/), or MATLAB. The sample sizes are shown in the figures. The minimum sample size of the electrophysiological experiment was set at 7 according to our previous power analysis (Ikeda et al., 2022). Statistical significance was set at *P* < 0.05. Full statistical reporting is shown in Table 1.

**Table 1.**
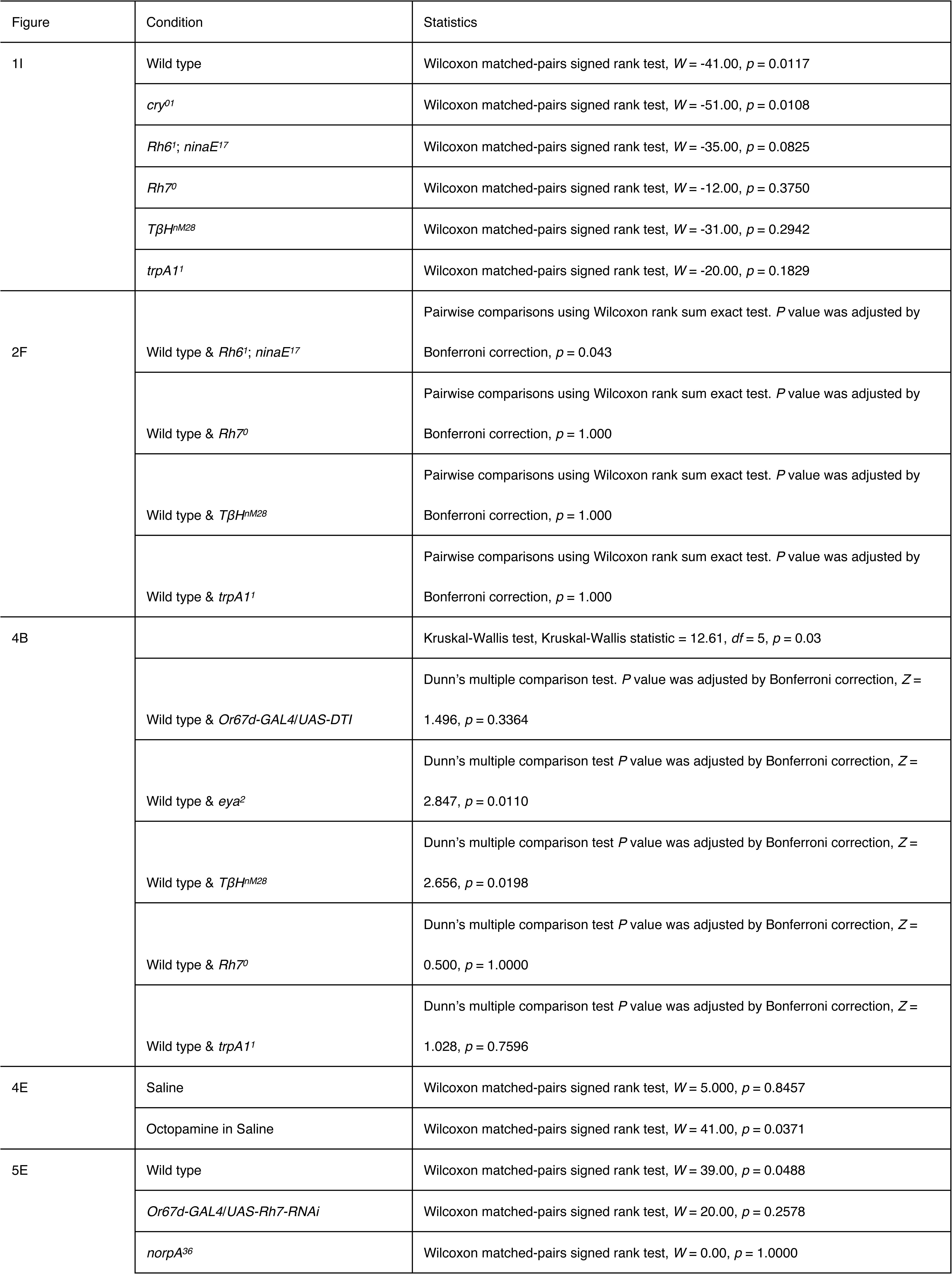
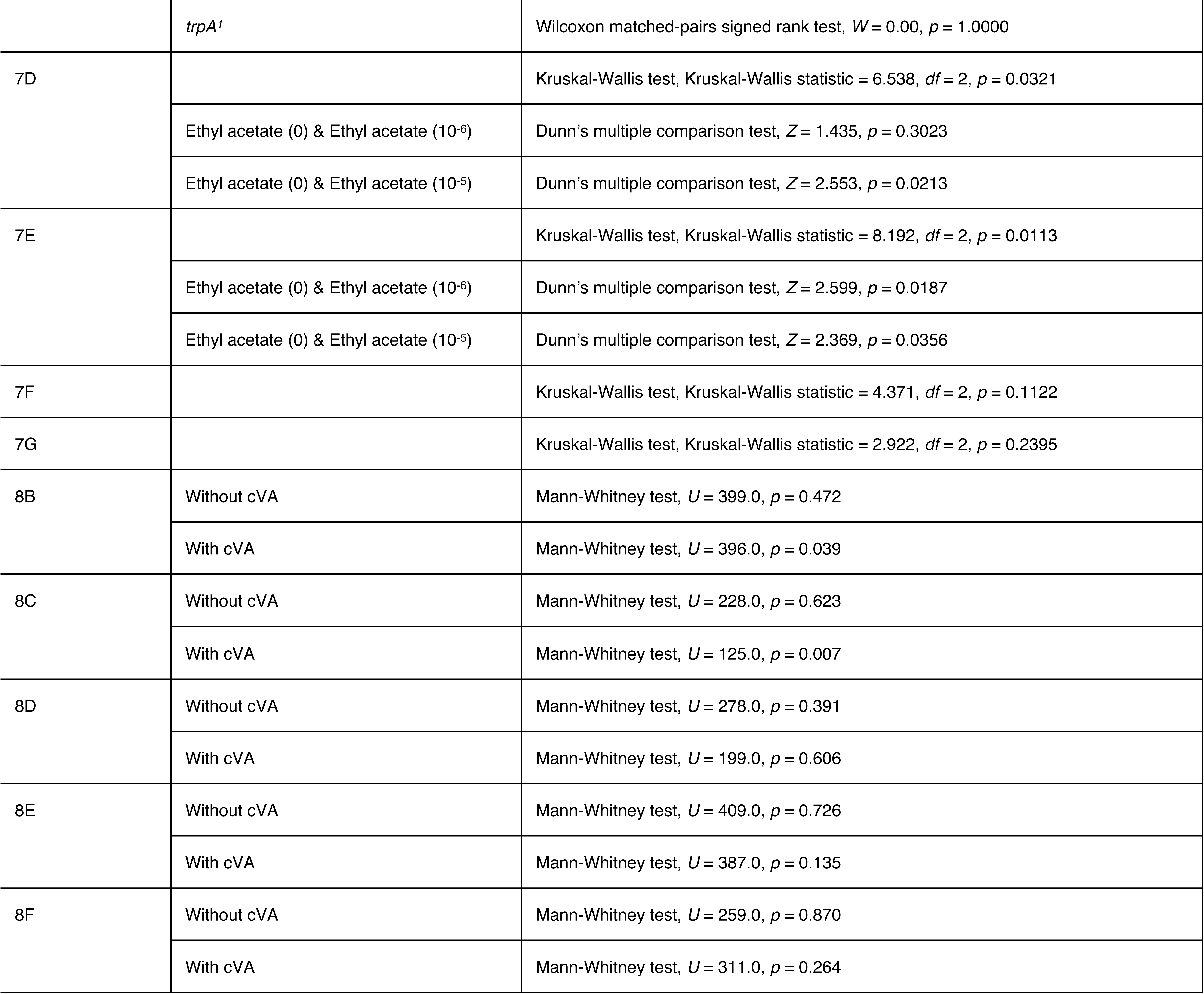
Detailed statistic values.

## Results

### Photoreceptor proteins indispensable for light-evoked firing rate increases in OSNs

We previously reported that transient blue light stimulation during odor stimulation increased the firing rates of OSNs expressing Or67d by approximately 2 Hz (Fig. 1A,B,C,I), which was mediated by the change in extracellular field potential around retinal photoreceptor cells (Ikeda et al., 2022). The Or67d-expressing OSN is the sole OSN housed in the T1 sensillum, the largest spine-shaped sensillum in the proximal part of the antenna (Shanbhag et al., 1999; Fig. 1A), and responds specifically to the male pheromone *cis*-vaccenyl acetate (cVA) (Clyne et al., 1997; van Naters and Carlson, 2007).

In this study, we investigated which photoreceptor proteins are indispensable for light-evoked increases in the firing rate in the OSNs. In *Drosophila*, *cryptochrome* (*cry*) and *rhodopsin* (*Rh1*, *5*, *6*, and *7*) have been identified as blue light receptor genes (Senthilan and Helfrich-Förster, 2016). *Rh1*, *5*, and *6* are strongly expressed in the retinal photoreceptor cells, whereas *cry* and *Rh7* are highly expressed in the brain (Senthilan and Helfrich-Förster, 2016; Ni et al., 2017). We observed a statistically significant light-evoked firing rate increase in *cry^01^* mutants but not in the double mutant *Rh6^1^*; *ninaE^17^*, which lacks expression of both *Rh6* and *Rh1* (Fig. 1D,E,I). The electroretinogram (ERG) response to blue light was significantly reduced in *Rh6^1^*; *nina^17^* mutants (Fig. 2A,B,F), and this result is consistent with our previous finding that light-evoked firing rate increases in the OSNs are mediated by the extracellular activity of photoreceptor cells in the compound eyes (Ikeda et al., 2022). In contrast, light-evoked firing rate increases in the OSNs were not observed in *Rh7^0^* mutants, although responses induced by cVA stimulation were normal (Fig. 1F). The ERG responses to blue light in *Rh7^0^* mutants were comparable to those in wild type flies (Ni et al., 2017; Fig. 2C,F). We previously reported that the light-evoked firing rate increases were not observed in flies whose heads, except for the antenna, were covered with black paint (Ikeda et al., 2022), indicating that blue light reception by Rh7 within the antenna does not contribute to the light-evoked firing rate increases. These results suggested that the functional role of Rh7 is to mediate the firing rate increase in the OSNs when the light information is relayed to the antenna.

**Figure 2.**
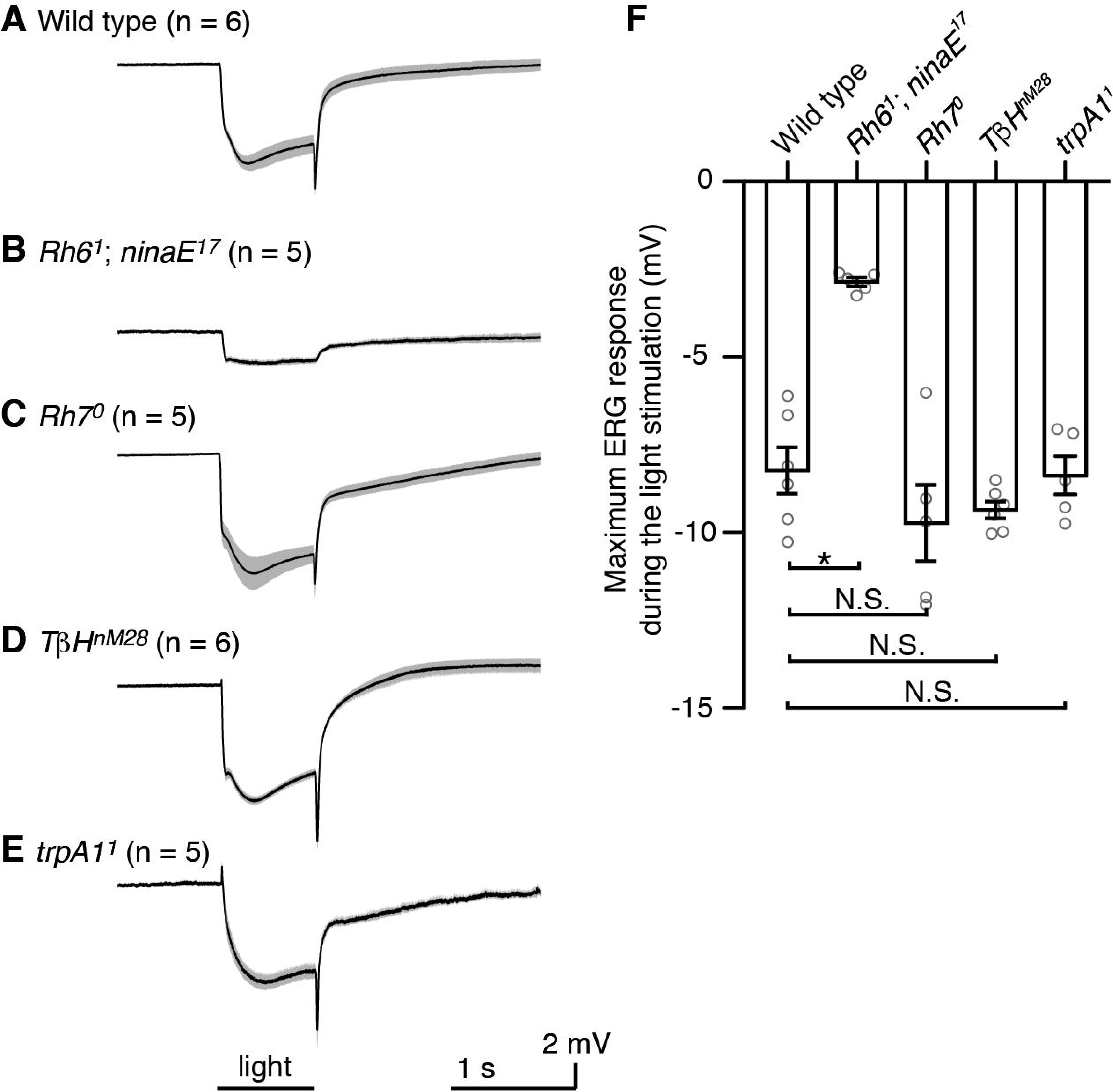
Electroretinogram recording. ***A-E,*** Electroretinogram (ERG) recorded from a compound eye in wild type (***A***), *Rh6^1^*; *nina^17^* (***B***), *Rh7^0^*(***C***), *TβH^nM28^* (***D***), *trpA1^1^* (***E***) mutants. ***F***, Maximum amplitudes of ERG deflection during the light stimulation. Shading and error bars represent the standard errors of the mean. *p* values were adjusted by Bonferroni correction after pairwise comparisons using Wilcoxon rank sum exact test. *, *p* < 0.05; N.S., not statistically significant. See Table 1 for full statistical reporting.

### Octopaminergic neurons relay the light information through the antennal nerve to induce light-evoked firing rate increases in the OSNs

How, then, is the visual information relayed from the retinal photoreceptor cells to the antennal OSNs? Since amputation of the antennal nerve abolished the light-evoked firing rate increases (Ikeda et al., 2022), we screened efferent neurons within the antennal nerve that connect the antenna and brain. In addition to sensory neurons, aminergic efferent fibers in the antennal nerve have been identified in various insect species (Busch et al., 2009; Pauls et al., 2018; Siju et al., 2008; Watanabe et al., 2014). Therefore, we labeled octopaminergic, serotonergic, histaminergic, and dopaminergic neurons using GAL4 lines (*NP7088* and *Tdc2-GAL4*, Busch et al., 2009 and Cole et al., 2005; *Hn.493-* and *Hn.819-GAL4*, Lee et al., 2011; *Ddc-*[*HL7-71* and *HL9-61*] and *TH-GAL4*, Claridge-Chang et al., 2009 and Ueno et al., 2012) and commercial antibodies and confirmed that the antennal nerve in *Drosophila* contains octopaminergic neurons but no other aminergic neurons (Fig. 3A; Busch et al., 2009; Pauls et al., 2018). Octopamine is a monoamine that is structurally related to norepinephrine in vertebrates. Detailed analyses of the projection pattern of the octopaminergic neurons within the antennae revealed a few fibers innervated the medial and lateral side of the third antennal segment housing the olfactory sensilla, but they did not project into each T1 sensillum containing the Or67d-expressing OSN (Fig. 3B,C). To examine whether the octopaminergic neurons contribute to the light-evoked firing rate changes, we recorded the neural responses of the T1 sensillum in *tyramine* β*-hydroxylase* (*TβH^nM28^*) mutants, which lack octopamine (Certel et al., 2007). The Or67d-expressing OSNs in the *TβH^nM28^* mutants exhibited normal cVA responses but did not show light-evoked firing rate increases (Fig. 1G,I). However, ERG responses to blue light stimulation revealed photoreception in the retinae in *TβH^nM28^*mutants was normal (Fig. 2D,F), indicating that light information is relayed to the antenna via octopaminergic neurons.

**Figure 3.**
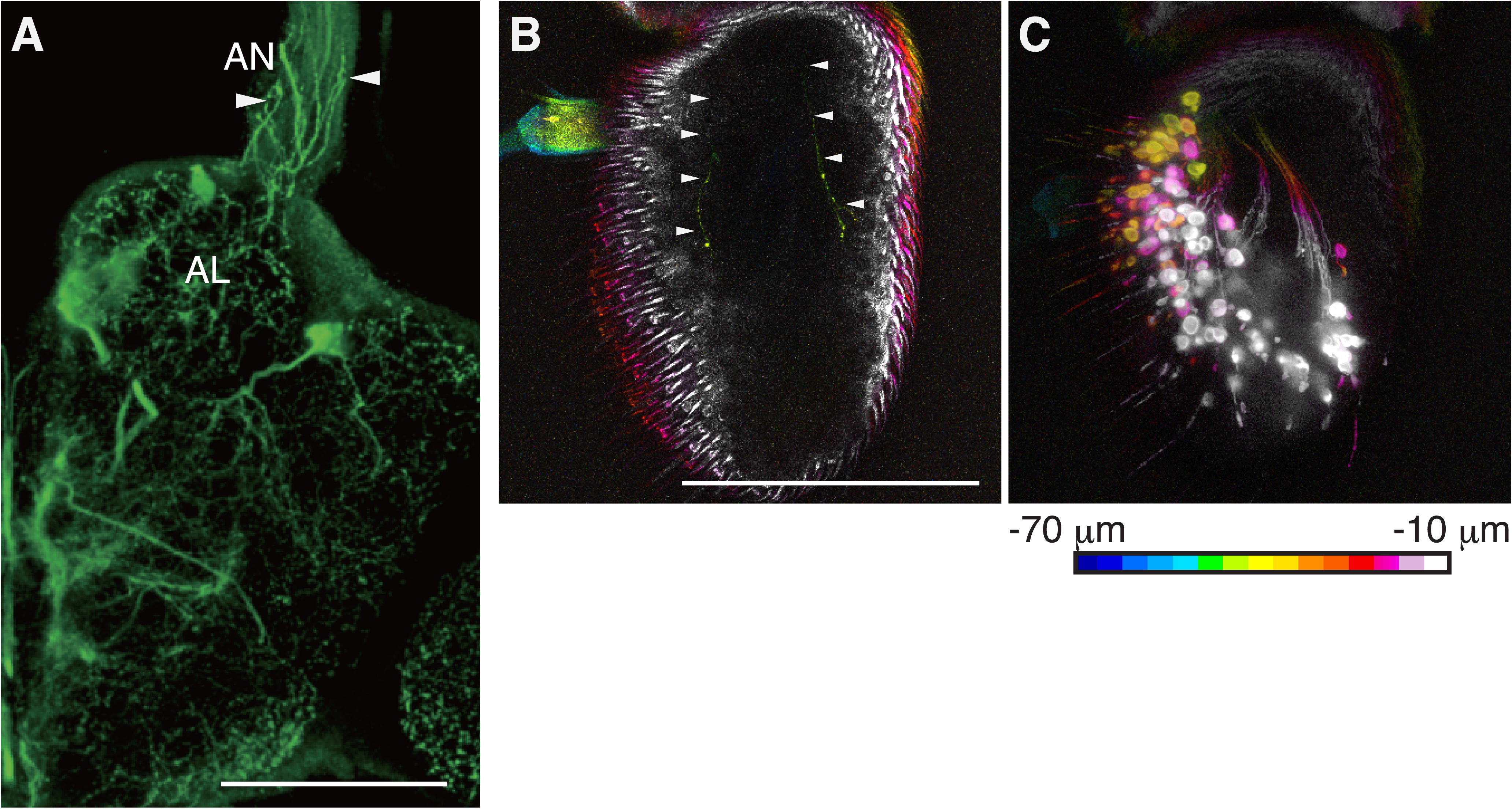
Octopamine application increased the firing rates of OSNs during odor stimulation. ***A,*** Efferent octopaminergic fibers, indicated by arrowheads, were labeled within the antennal nerve by *Tdc2-GAL4.* Anterior is to the top and lateral is to the right. AL, antennal lobe; AN, antennal nerve. ***B,C,*** The octopaminergic fibers, indicated by arrowheads in ***B***, do not terminate in close to each Or67d-expressing OSN cell body shown in ***C*** in the third antennal segment. The depth from the anteriormost surface of the antenna is represented by color as shown in the bottom. The signals of anterior surfaces of the antennae are not included in these images, as autofluorescence hides the signals of the octopaminergic fibers. It should be noted that the anterior surface does not include any signals of the octopaminergic fibers labeled by *Tdc2-GAL4.* Dorsal and medial are to the top and right, respectively. Scale bars = 100 μm. Genotypes: (***A,B***) *w*; *Tdc2-GAL4*/*UAS-GFP*; *UAS-20XmCD8::GFP*/*+*; (***C***) *w*; *Or67d-GAL4*; *10XUAS-mCD8::GFP*.

### Light-evoked increases in the field potential are mediated by octopaminergic neurons

The extracellular potential recorded from the sensillum reveals the local field potential (LFP) as well as OSN spiking during sensory stimulation. Interestingly, we found that light stimulation caused positive 1.1-mV LFP deflections in wild type flies on average (Fig. 4A,B) in contrast to the negative LFP deflections evoked by odor stimulation (Ayer and Carlson, 1992). This result was surprising, as increased excitation of the OSNs was previously found to correlate with decreased extracellular LFP (Zhang et al., 2019). Since the light increased the firing rate in the OSNs (Fig. 1C), we expected decreased rather than increased LFP deflections during the light stimulation. This result would imply that the light-evoked increase in LFP may be caused by cells other than the OSN within the sensillum. We thus investigated whether OSNs are indispensable for this light-evoked increase in LFP by expressing the temperature-sensitive *diphtheria toxin* (*DTI*) (Bellen et al., 1992; Han et al., 2000) in the Or67d-expressing OSNs to conditionally introduce cell death after eclosion (Fig. 4C,D). Extracellular potentials recorded from the T1 sensillum lacking the OSN revealed that light-evoked positive LFP deflections were comparable to those in wild-type flies, whereas we observed neither neural firings nor odor-driven negative LFP deflections (Fig. 4A,B). The T1 sensillum contains three types of auxiliary cells, namely the trichogen, tormogen, and thecogen cells, that surround the Or67d-expressing OSN and generate the transepithelial potential (Fig. 1A; Stengl, 2010). This suggests that the light-evoked increase in the LFP is primarily caused by the auxiliary cells in an OSN-independent manner.

**Figure 4.**
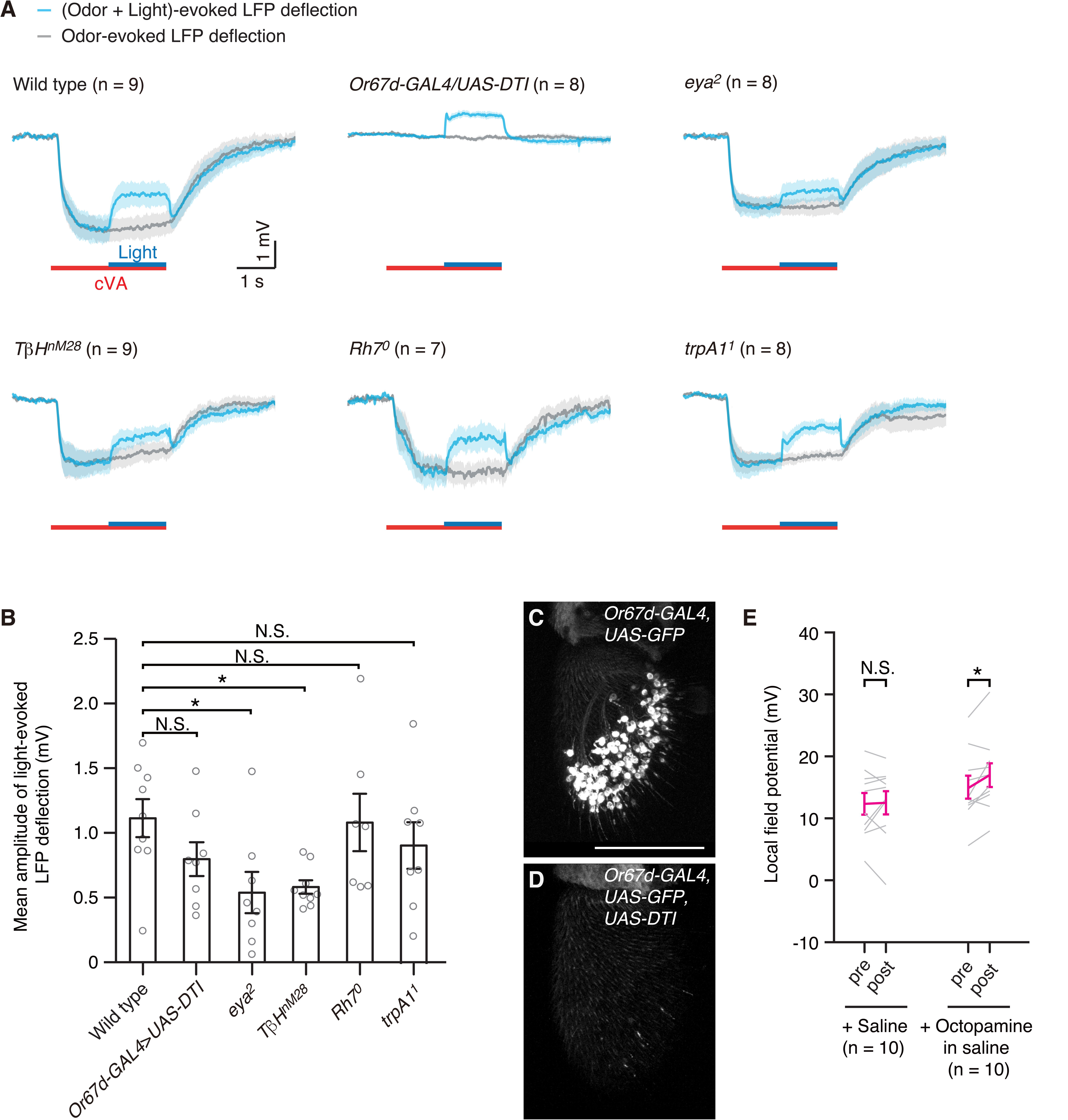
Local field potential deflections evoked by odor and light stimuli recorded from the T1 sensillum. ***A***, Local field potential (LFP) deflections evoked by odor (cVA) and light stimuli recorded from the T1 sensilla in wild type, *Or67d-GAL4*>*UAS-diphtheria toxin* (*DTI*), *eya^2^*, *TβH^nM28^*, *Rh7^0^*, and *trpA1^1^* flies. Gray represents the LFP responses to the only odor stimulation, whereas cyan shows responses to combined odor and light stimulation. Shading represents the standard error of the mean. ***B*,** Light-evoked LFP deflection amplitudes averaged during the light stimulation. ***C*,*D*,** OSNs labeled with GFP under *Or67d-GAL4* expression (***C***) were not observed in ***D***, as the *diphtheria toxin* gene was co-expressed to induce cell death. 3D-reconstructed third antennal segments are shown. Dorsal is to the top, lateral is to the right. Scale bar = 100 μm. ***E*,** The LFP in the T1 sensillum was upregulated by octopamine application into the hemolymph even in the absence of the Or67d-expressing OSNs. Each gray line shows the LFP recorded from a different animal and magenta represents the mean with the standard error of the mean. Genotypes: (***C***) *w*; *Or67d-GAL4*/*UAS-GFP*; *+*/*20XUAS-mCD8::GFP*; (***D***) *w*; *Or67d-GAL4*/*+*; *UAS-DTI*/*20XUAS-mCD8::GFP*; (***E***) *w*; *Or67d-GAL4*/*+*; *UAS-DTI*/*+*. Error bars indicate the standard errors of the mean. For statistical analyses, Kruskal-Wallis test followed by Dunn’s multiple comparison test (***B***) and Wilcoxon matched-pairs signed rank tests (***E***) were performed. *, *p* < 0.05; N.S., not statistically significant. See Table 1 for full statistical reporting.

The amplitude of the light-evoked LFP deflection within the antennal sensillum was significantly smaller in *eyes absent*^2^ (*eya^2^*) mutants, which lack compound eyes (Choi and Benzer, 1994), than in wild type flies, indicating that the positive deflection was primarily caused by photoreception in the compound eyes (Fig. 4A,B). The small amplitude of the light-evoked LFP deflection in *eya^2^* was comparable to that in *TβH^nM28^* (Fig. 4A,B), consistent with our finding that light information is relayed to the antenna by octopaminergic neurons. The light-evoked firing rate increases in the OSNs were not observed in *eya^2^*as well as *TβH^nM28^* (Ikeda et al., 2002; Fig. 1G), implying that the amplitude of the LFP deflection in these mutants would not be sufficient to cause the firing rate increases.

To reveal whether octopamine is responsible for regulating the LFP, we created an opening in the thorax in the flies lacking the Or67d-expressing OSNs and covered it with saline to make the hemolymph continuous with the saline. We then applied octopamine to the saline (200 μM, in saline), allowing three minutes for it to reach the antennae by diffusion. Whereas we did not observe a statistically significant increase in the LFP in control where saline only was applied, saline containing octopamine upregulated the LFP by 1.9 mV in the T1 sensillum (Fig. 4E). We thus concluded that the octopaminergic neurons relay light information to the antenna and their output generates the light-evoked positive LFP deflections in the sensillum lymph by acting on the auxiliary cells.

### Positive deflection in the extracellular field potential causes the firing rate increases in the OSNs

Since the octopaminergic neurons upregulate the LFP (Fig. 4A,E) and their output is indispensable for the light-evoked firing rate increases in the OSNs (Fig. 1G), we next investigated whether the positive LFP deflection is the cause of the firing rate increases during light stimulation. We injected a 5-pA current into the lymph of a T1 sensillum during the odor stimulation. The current caused 1.55 ± 0.13 mV positive LFP deflections in wild type, which significantly increased the firing rates of the Or67d-expressing OSNs by 2.1 Hz on average (Fig. 5A,E), comparable to the light-evoked firing rate increases. This result indicates that the light-evoked firing rate increases in the OSNs were caused by the positive deflection of the LFP.

**Figure 5.**
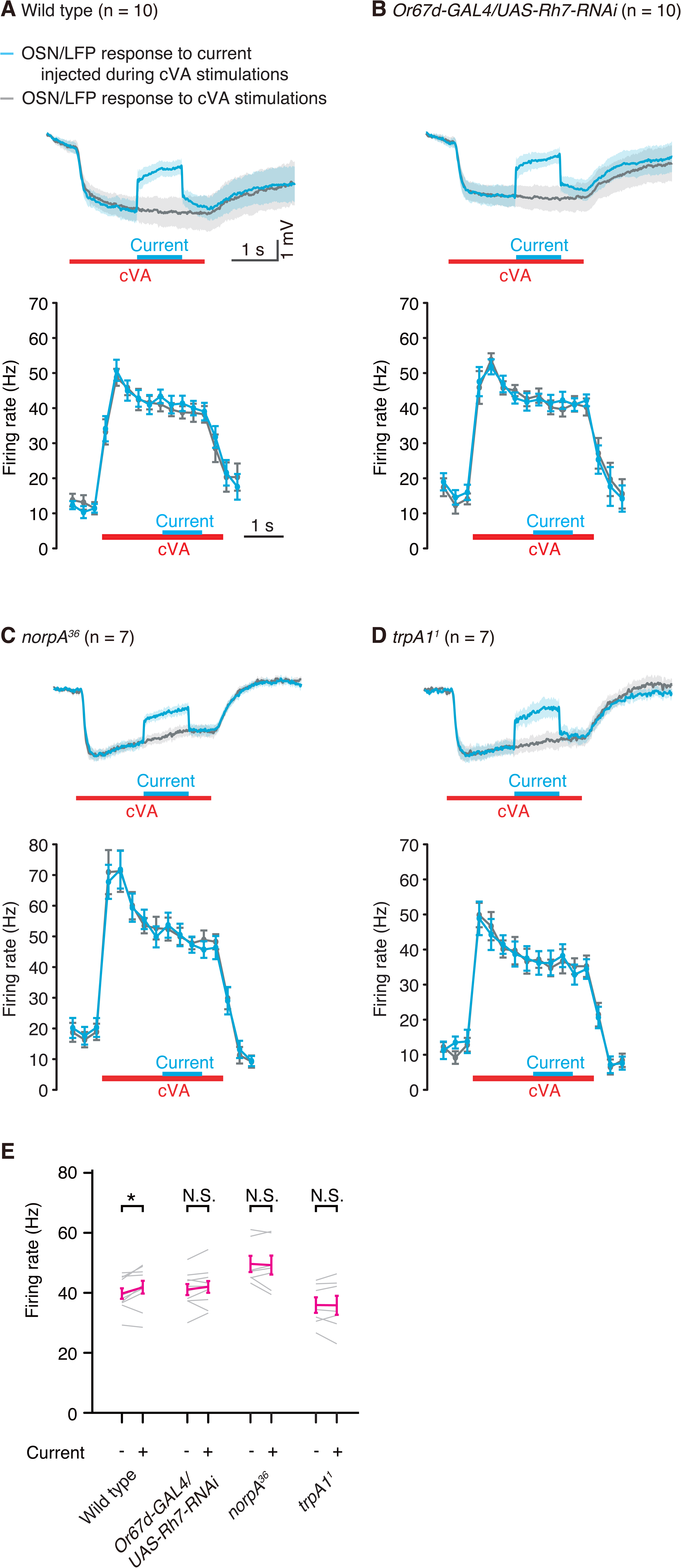
Positive LFP deflection induced by current injection increased the firing rates of Or67d-expressing OSNs during cVA stimulation. A current transiently injected into the T1 sensillum during cVA stimulation evoked positive LFP deflections in wild type (*A*, top), *Or-67d-GAL4*/*UAS-Rh7-RNAi* flies (*B*, top), *norpA^36^*(*C*, top), and *trpA1^1^* (*D*, top), whereas the current injection resulted in the increase in the firing rates of Or67d-expressing OSNs only in wild type (*A*, bottom), but not in *Or67d-GAL4*/*UAS-Rh7-RNAi* (*B*, bottom), *norpA^36^* (*C*, bottom), and *trpA1^1^* (*D*, bottom). Gray represents the LFP responses to the sole cVA stimulation (top) and firing rates of the OSNs (bottom), whereas cyan shows LFP responses to current injection applied during the cVA stimulation (top) and firing rates of the neurons (bottom). Shading represents the standard error of the mean. (*E*) The firing rates of Or67d OSNs were increased significantly in wild type, but not in *Or-67d-GAL4*/*UAS-Rh7-RNAi*, *norpA^36^*, and *trpA1^1^* by current injection. For statistical analyses, Wilcoxon matched-pairs signed rank tests were performed. *, *p* < 0.05; N.S., not statistically significant. See Table 1 for full statistical reporting.

Interestingly, the light-evoked positive LFP deflection in *Rh7^0^*mutant flies was comparable to that of wild type (Fig. 4A,B), although the OSNs in *Rh7^0^* did not show light-evoked firing rate increases (Fig. 1F). This suggested that OSNs lacking *Rh7* expression might be dysfunctional in responding to the LFP changes. We thus injected a 5-pA current into the T1 sensillum in which *Rh7* expression was downregulated by RNAi specifically in the Or67d-expressing OSN during the odor stimulation to artificially evoke a positive LFP deflection. Although the current injection evoked 1.42 ± 0.11 mV positive LFP deflections, we did not observe a significant increase in the firing rates of the OSNs (Fig. 5B,E), indicating that Rh7 is indispensable for regulating the firing rates of the OSNs in response to the LFP changes.

Rh7 functions as a detector of bitter compounds in a light-independent manner and initiates signaling cascade including phospholipase C-β (PLC-β) and transient receptor potential channel A1 (TRPA1) (Leung et al., 2020). We therefore investigated whether the positive LFP deflection evoked the firing rate increases of the Or67d-expressing OSNs in PLC-β (*norpA^36^*) and *trpA1^1^* mutants. Although the current injection induced 1.12 ± 0.05 and 1.37 ± 0.10 mV positive LFP deflections in *norpA^36^* and *trpA1^1^* mutants, respectively, we did not observe a significant increase in the firing rates of the OSNs (Fig. 5C,D,E). Consistent to these results, the light-evoked firing rate increases during odor stimulation were not observed in the *trpA1^1^* mutant (Fig. 1H,I), though the *trpA1^1^* mutant showed normal ERG responses comparable to wild type (Fig. 2E,F). These results suggest that Rh7 changes the firing rates of the OSNs by opening TRPA1 channels.

### Structural analysis of Rh7

To our knowledge, the entire protein structure of Rh7 has not been reported. In the present study, the predicted structure of Rh7, generated using AlphaFold2, has seven transmembrane helices (helices 1 to 7) with an additional three helices (helices 8 to 10) located in the membrane-extrinsic, intracellular region (Fig. 6A). Helices 8 and 10 lie parallel to the membrane surface, whereas helix 9 is perpendicular to the transmembrane. Notably, helices 1 to 8 are highly conserved, and helix 9 is partially conserved in the predicted rhodopsin 1 (Rh1) structure (Fig. 6B and Supplementary Fig. 1), making the presence of the additional C-terminal region with helix 10 remarkable for Rh7. Indeed, it has been observed that the C-terminus of Rh7 is twice as long as those in other rhodopsins (Senthilan and Helfrich-Förster, 2016). This structural divergence suggests a potential functional role may exist for the extended region of Rh7 in sensing and responding to extracellular electric fields.

**Figure 6.**
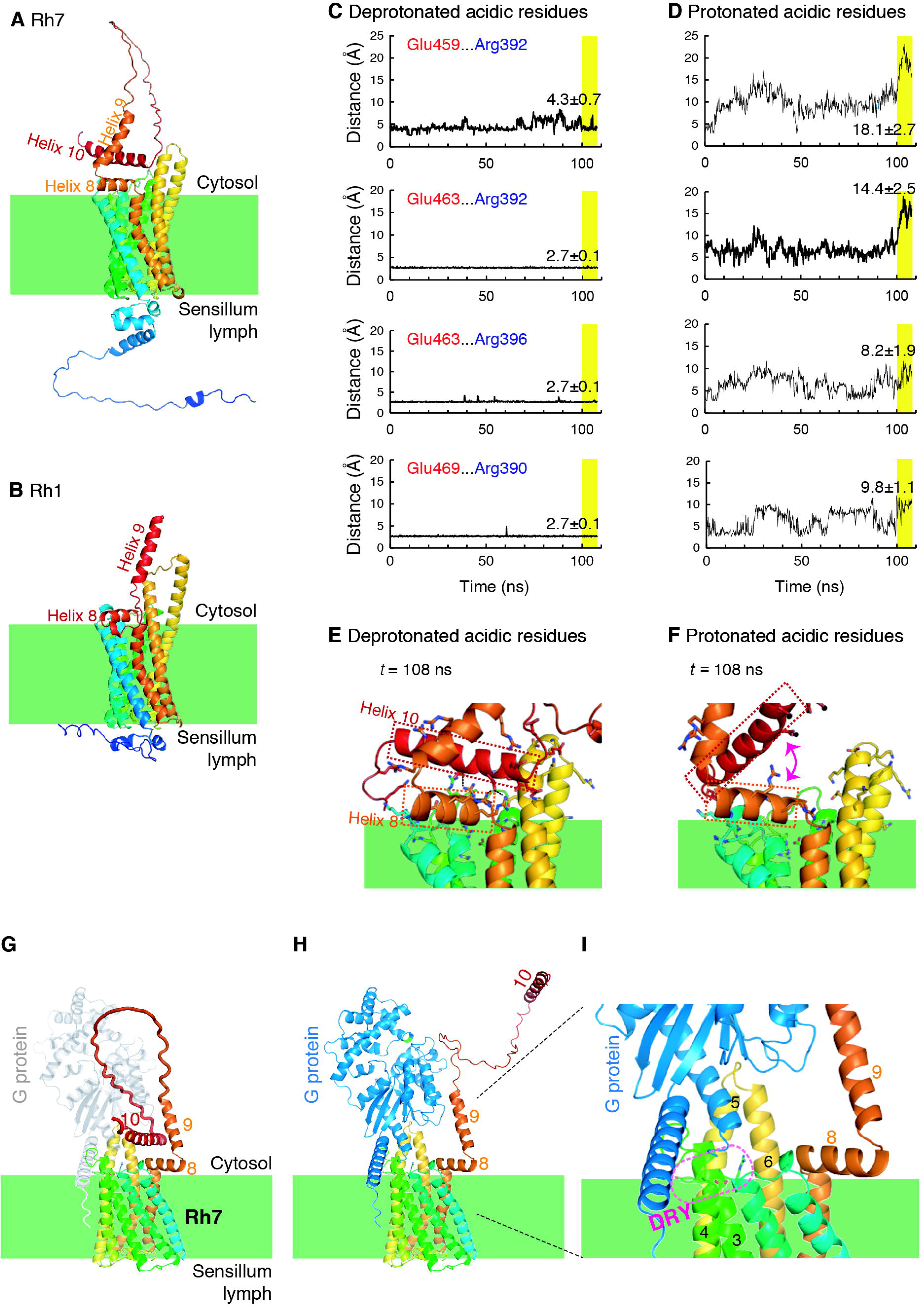
Predicted rhodopsin 7 (Rh7) structures. ***A*,*B*,** Comparison of predicted structures of Rh7 (***A***) and Rh1 (***B***). ***C*-*F*,** Molecular dynamics (MD) simulations for Rh7. (***C***) H-bond distances (Å) between helices 8 and 10 in the standard protonation state of the acidic residues in helix 10. (***D***) H-bond distances (Å) between helices 8 and 10 upon protonation of the acidic residues in helix 10. The average and standard deviations of the distances were obtained during the period from 100 to 108 ns (highlighted in yellow). (***E***) Resulting geometry for Rh7 after a 108-ns MD run in the standard protonation state of the acidic residues in helix 10. (***F***) Resulting geometry for Rh7 after a 108-ns MD run upon protonation of the acidic residues in helix 10. Helices 8 and 10 are surrounded by orange and red dotted lines, respectively. Magenta double arrow indicates the dissociation of helix 10 from helix 8. ***G*-*I*,** Modeled Rh7/G-protein complex structure. (***G***) G-protein unbound conformation of Rh7, where helix 10 interacts with helix 8. G-protein (gray) is shown to represent the steric hindrance with the long loop connecting helices 9 and 10. (***H***) G-protein bound conformation of Rh7, where helix 10 is separated from helix 8 to facilitate binding of the G-protein. (***I***) Binding interface in the Rh7/G-protein complex. Magenta dotted oval indicates the DRY motif (Asp212, Arg213, and Tyr214), which plays a crucial role in G-protein binding (Franke et al., 1992). Each number in ***G***-***I*** indicates the helix number. See Supplementary Fig. 1 and 2 for additional information regarding Rh7, Rh1, and G-protein structures.

The electrical potential across the membrane induces an accumulation of negative charges in the intracellular domain during the positive LFP deflection, which drives the uptake of positively charged protons (Casey et al., 2010; Alvares et a., 2021). Consequently, this polarization potentially triggers local protonation of titratable residues, particularly acidic residues, on the intracellular surface of the protein.

Of particular interest, helix 10 in Rh7 harbors a cluster of acidic residues, including Glu459, 460, 462, 463, and 469 (Supplementary Fig. 1). Many of these residues are likely involved in hydrogen bond (H-bond) interactions with basic residues such as Arg in helix 8. To elucidate further details of the interactions between helices 8 and 10 in equilibrium, we conducted molecular dynamics (MD) simulations.

In the standard protonation state, where acidic residues remain deprotonated and basic residues are protonated, stable H-bonds were observed between helices 10 and 8. Four H-bonds, namely Glu459…Arg392, Glu463…Arg392, Glu463…Arg396, and Glu469…Arg390 between helices 10…8, remained stable during MD simulations (Fig. 6C,E). However, upon protonation of the acidic residues in helix 10, these H-bonds became unstable, resulting in their loss and the subsequent release of helix 10 from helix 8 (Fig. 6D,F). These observations suggest that applying a positive LFP deflection may induce a significant conformational change in Rh7, leading to the dissociation of helix 10 from helix 8 and the associated transmembrane region.

To further elucidate the role of helix 10 in Rh7, we modeled the complex structure with G-protein based on the reported cryo-EM structure of the rhodopsin/G-protein complex (Gao et al., 2019) (Supplementary Fig. 2). The modeled Rh7/G-protein complex structure reveals that the binding of G-protein to Rh7 is inhibited by the presence of the long loop connecting helices 9 and 10 (Met416 to Thr458), as long as helix 10 interacts with helix 8 in the standard protonation state (Fig. 6G). In contrast, upon protonation of its acidic residues, helix 10 dissociates from helix 8. Consequently, the binding of G-protein to Rh7 appears to be facilitated, as the long loop no longer occupies the binding space surrounding the DRY motif (Asp212, Arg213, Tyr214) required for G-protein interaction (Franke et al., 1992; Fig. 6H,I). These observations suggest that the release of helix 10 and the accompanying long loop are likely prerequisites for G-protein binding at Rh7.

### Rh7 contributes to ephaptic inhibition between OSNs

Su et al. (2012) revealed ephaptic inhibition between OSNs within the same sensillum, where excitation of one neuron (A) caused a negative LFP deflection, resulting in inhibition of the other neurons (B) via the electric field effect. As Rh7 is indispensable for firing rate increases in the OSNs in response to the positive LFP deflection that changes the electric field, we investigated if Rh7 contributes to the ephaptic inhibition. We recorded the odor responses from the ab2 sensilla, which contains two OSNs, ab2A and ab2B, that express olfactory receptors Or59b and Or85b, respectively (Fig. 7A; Hallem et al., 2004). These two OSNs can be readily distinguished by spike amplitudes; larger and smaller spikes originate from ab2A and ab2B, respectively. Ethyl acetate specifically excites ab2A, whereas hexanol induces spikes in ab2B (Fig. 7B; Bruyne et al., 2001). When hexanol odor was transiently applied during sustained stimulation with ethyl acetate, the odor response of ab2B to hexanol was inhibited by excitation of ab2A in an ethyl acetate concentration-dependent manner in wild type flies (Fig. 7C,D). The firing rates of ab2B in response to hexanol (10^0^) was maximal when a clean air puff was applied in place of ethyl acetate, whereas the firing rates gradually decreased when the concentration of ethyl acetate in the flask was increased from 10^-6^ to 10^-5^ (Fig. 7D). The concentrations indicated are those of the solutions prepared in the flask and vial, from which saturated vapor was applied to the fly antennae. This ephaptic inhibition was not dependent on light, as we observed the inhibition even in darkness (Fig. 7E). In contrast to wild type, we did not observe ephaptic inhibition in *Rh7^0^* mutants (Fig. 7F,G). Because the firing rates of ab2B in *Rh7^0^* were larger in response to hexanol (10^0^) than those in wild type (Fig. 7F), we also analyzed the firing rates of ab2B in response to a lower concentration of hexanol (10^-3^) during sustained stimulation with ethyl acetate (Fig. 7G). In wild type, ab2B response to the lower concentration of hexanol (10^-3^) was also inhibited during sustained stimulation with ethyl acetate (Supplementary Fig. 3A). At these hexanol concentrations, we did not observe significant ephaptic inhibition in ab2B in *Rh7^0^* in spite of the fact that the firing rates of ab2A and the amplitudes of LFP deflections during ethyl acetate stimulation in *Rh7^0^* were comparable to those in wild type (Supplementary Fig. 3B,C). We thus concluded that Rh7 mediates the firing rate changes in OSNs in response to the LFP change, which is not dependent on light.

**Figure 7.**
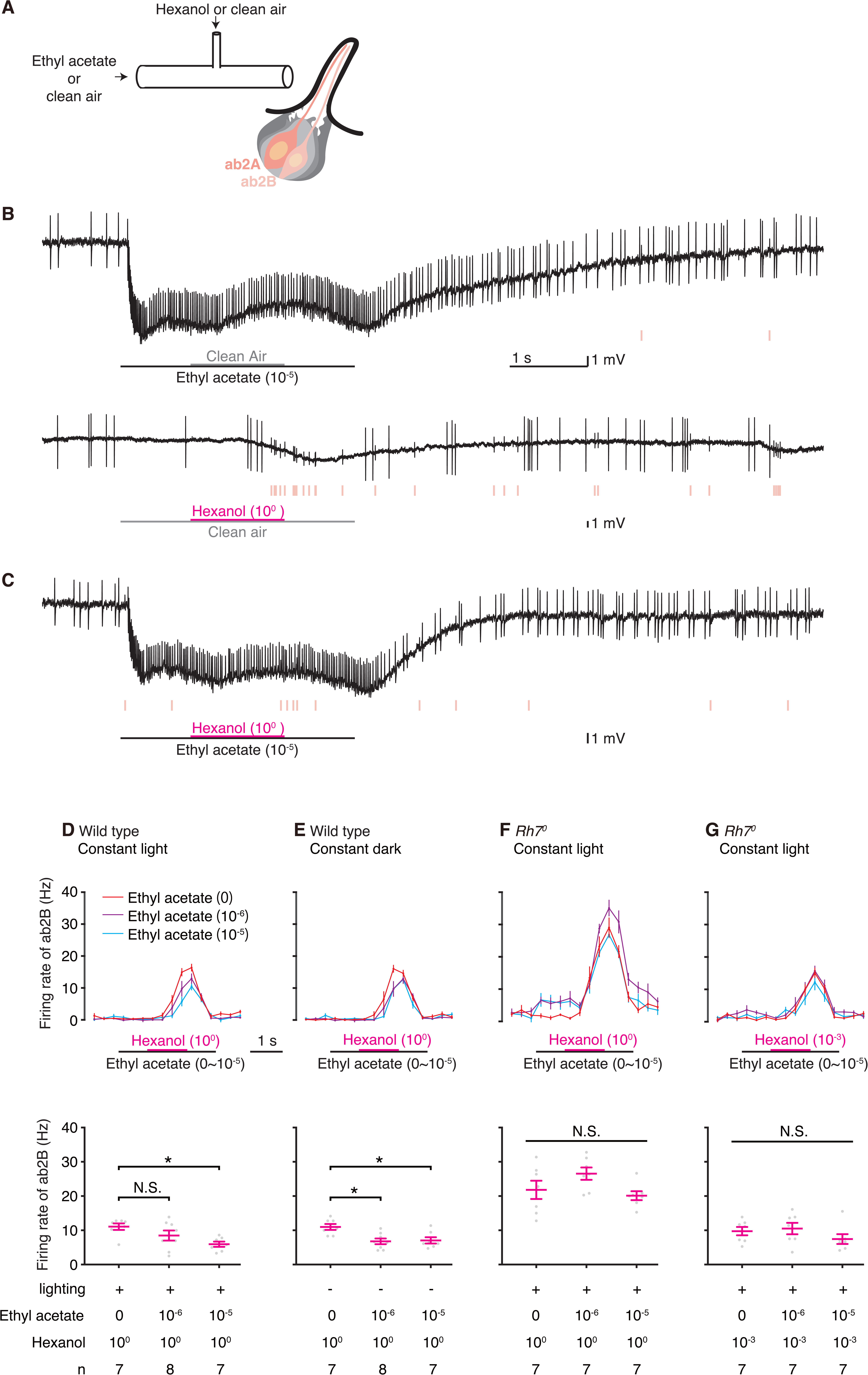
Rhodopsin 7 contributes to ephaptic inhibition between OSNs within a sensillum. ***A***, Air passed through a vial containing hexanol or an open vial (clean air) joins air passed through a flask containing ethyl acetate diluted in paraffin oil or an open flask (clean air) and stimulates the ab2 sensillum in the fly antenna. The ab2 sensillum houses two olfactory sensory neurons, ab2A and ab2B as well as auxiliary cells, colored in gray. ***B*,** Extracellular recording from the ab2 sensillum showed spikes of two amplitudes in addition to LFP deflections. ab2A, represented by larger spikes, responded to ethyl acetate but not to hexanol, whereas ab2B, with smaller spikes (indicated by pink bars below traces), responded to hexanol but not to ethyl acetate. It should be noted that the concentrations of the chemicals shown here are those prepared in the flask or vial and that evaporated vapor in the flask or vial was applied to the fly. Horizontal bars represent the timing each air puff was applied to the tube. Vertical black bars, 1mV. ***C*,** Air puff containing ethyl acetate slightly inhibited the ab2B responses to hexanol. ***D-G*,** The response of ab2B to hexanol was inhibited by the presence of ethyl acetate in a concentration-dependent manner in wild type in both constant light (***D***) and dark (***E***) conditions, but not in the *Rh7^0^* mutants (***F*,*G***). Top panels show the firing rates in each bin of 300 ms. Magenta bars represent 1.2 s of hexanol puff, whereas black bars indicate the timing of 3 s puff of three different concentrations of ethyl acetate (0-10^-5^). Error bar indicates the standard error of the mean. Bottom graphs show the mean firing rates of ab2B during the period in which it showed responses to hexanol (for 1.2 s from 0.6 s after the onset of hexanol puffs). Each gray dot indicates the firing rate of a different animal, and magenta represents the mean ± the standard error of the mean. For statistical analyses, Kruskal-Wallis tests followed by Dunn’s multiple comparisons were performed. *, *p* < 0.05; N.S., not statistically significant. See Table 1 for full statistical reporting and Supplementary Fig. 3 for the response of ab2B to hexanol (10^-3^) during sustained stimulation with ethyl acetate in wild type and ab2A and LFP responses to ethyl acetate.

### Behavioral response to cVA is modulated by light

Our study revealed that the blue light stimulation changed the firing rate of each Or67d-expressing OSN by approximately 2 Hz during cVA stimulation. cVA induces aggregation behavior in females (Bartelt et al., 1985). We next examined whether such a small change in the firing rate in each OSN can affect fly behavior. We compared the cVA responses between flies that showed the light-evoked firing rate increases during cVA stimulation and those that did not by investigating whether light stimulation changed the direction of fly movement. We prepared a double-layered chamber (Tanaka et al., 2021) and placed a virgin female in the upper cell of the chamber, above the solvent or cVA diluted in solvent located in the center of the lower cell. The female fly was separated from the odor source by a mesh barrier. We measured cos(*θ*) immediately after the offset of the light stimulation, where *θ* was the angle between the direction of fly movement and the vector joining the fly to the center of the cell at the light offset (Fig. 8A). When a wild-type female walked over the solvent, light stimulation did not cause a significant change in the direction of movement (Fig. 8B left), whereas fly movement became more oriented toward the center by light when cVA was placed in the lower cell (Fig. 8B right). Flies expressing tetanus toxin in the photoreceptor cells in the retinae, eyelets, and ocelli (*longGMR-GAL4*/*UAS-TNT*), in which light-evoked firing rate increases in the OSNs were normally observed (Ikeda et al., 2022), also showed significant changes in walking direction when cVA and light were paired (Fig. 8C). This result indicates that the light-evoked cVA-dependent change in the direction of movement was induced without synaptic relay of light information from the external photoreceptor cells. On the other hand, we did not observe significant light-evoked cVA-dependent changes in the orientation of fly movement in *eya^2^*and *TβH^nM28^* mutant flies (Fig. 8D,E). *TβH^nM28^*flies showed normal cVA responses and ERG responses to blue light, but not light-evoked firing rate increases in the OSNs (Figs. 1G,2D). In addition, downregulation of *Rh7* expression in the Or67d-expressing OSNs also impaired the light-dependent changes in the cVA response (Fig. 8F), demonstrating that Rh7 expression in the OSNs contributes to changes in the behavioral responses as well as neural responses to the odor in a light-context dependent manner. These results indicate that the light-evoked LFP deflections associated with light-evoked firing rate increases in the OSNs are the cause of the light-evoked behavioral responses to cVA.

**Figure 8.**
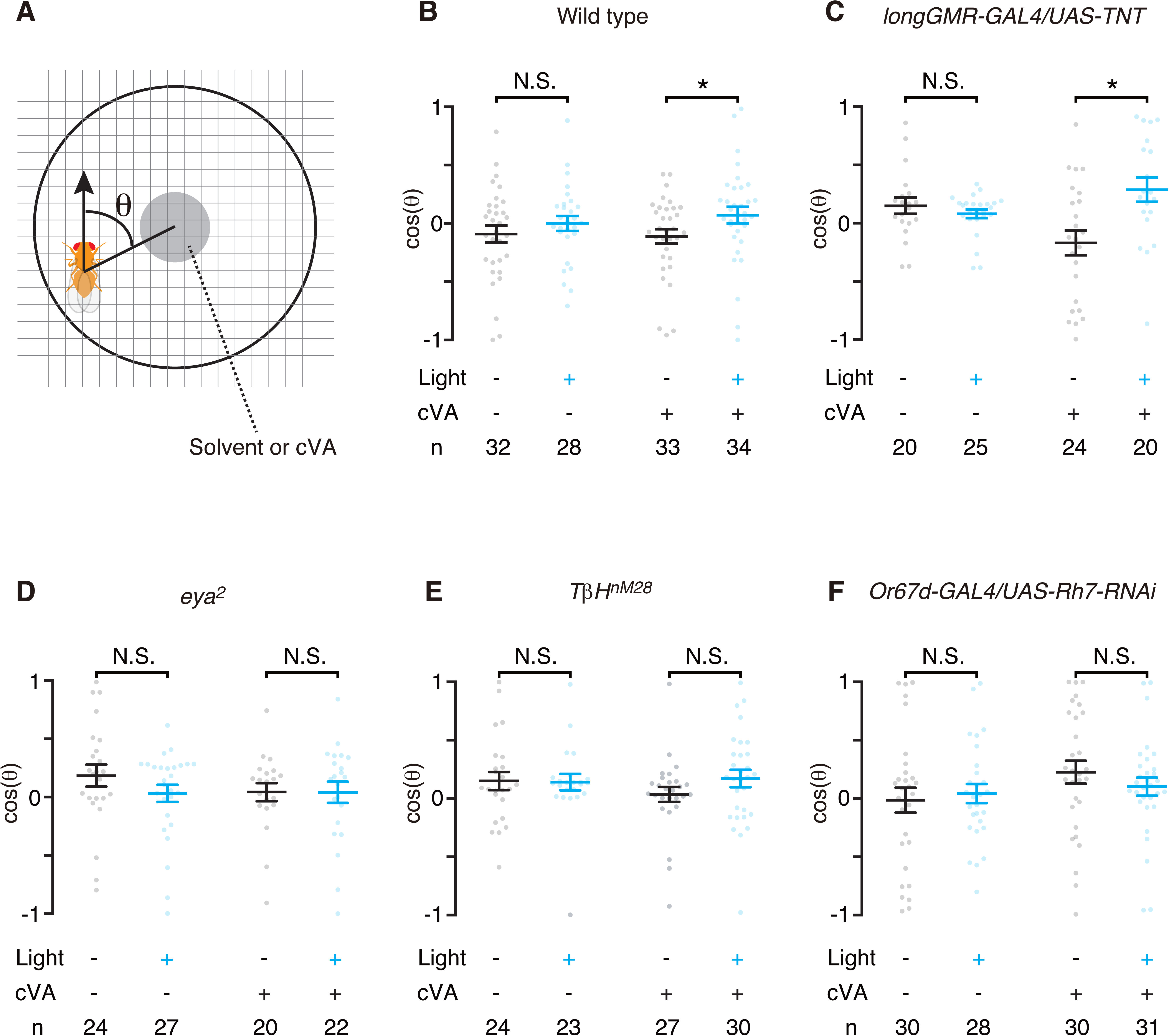
Behavioral responses to *cis*-vaccenyl acetate (cVA) were altered by simultaneous light stimulation. ***A***, Experimental paradigm. A female fly was placed on a mesh set above a filter paper containing 18 ng cVA or solvent and was video-recorded to analyze the walking direction. *θ* is the angle between the direction of the fly’s movement (arrow) and the vector that joins the centroid of the fly’s body to the center of the cell (solid black line) just after the offset time of the light stimulations. The fly moved toward the center of the cell at the offset time, when cos(*θ*) was 1. ***B-F*,** Light stimulation increased the value of cos(*θ*) significantly in wild type (***B***) and the flies expressing tetanus toxin (TNT) in external photoreceptor cells (*longGMR-GAL4*/*UAS-TNT*) (***C***) in the presence of cVA, which was not observed in the *eya^2^*(***D***), *TβH^nM28^* (***E***), and *Or67d-GAL4*/*UAS-Rh7-RNAi* (***F***). Each dot represents a single animal. Error bars indicate the standard errors of the mean. Mann-Whitney tests were performed for statistical analyses. *, *p* < 0.05; N.S., not statistically significant. See Table 1 for full statistical reporting.

## Discussion

Recent studies reported that rhodopsins contribute to thermosensation, hearing, proprioception, and gustation in a light-independent manner (Shen et al., 2011; Senthilan et al., 2012; Sokabe et al., 2016; Zaini et al., 2018; Leung et al., 2020). Similarly, we showed that light reception by Rh7 in the antenna does not contribute to the light-evoked firing rate increases of the OSNs, but rather, its functional role is to mediate responses to the LFP deflection within the olfactory sensillum (Fig. 9). The light-evoked LFP deflection is caused by the output of the octopaminergic neurons that acts on non-neuronal cells (Fig. 4). Rh7 and its downstream PLC-β and TRPA1 expressed in the OSN are indispensable for the firing rate changes in response to an externally imposed extracellular LFP deflection (Fig. 5). Moreover, Rh7 contributes to naive ephaptic inhibition where excitation of one OSN induces inhibition in another OSN by the electric field effect during the odor stimulation (Fig. 7). Conformational analyses of Rh7 suggest the helices 8 and 10 form H-bonds that, together with the long loop region, function as a voltage-dependent gate for G protein binding (Fig. 6). Finally, we showed that the firing rate increases in the OSNs evoked by the LFP deflection contribute to olfactory-visual integration (Fig. 8). These findings demonstrate that Rh7 regulates neural activity in response to continuous changes in extracellular electrical activity, thereby modulating behavioral patterns in a context-dependent manner.

**Figure 9.**
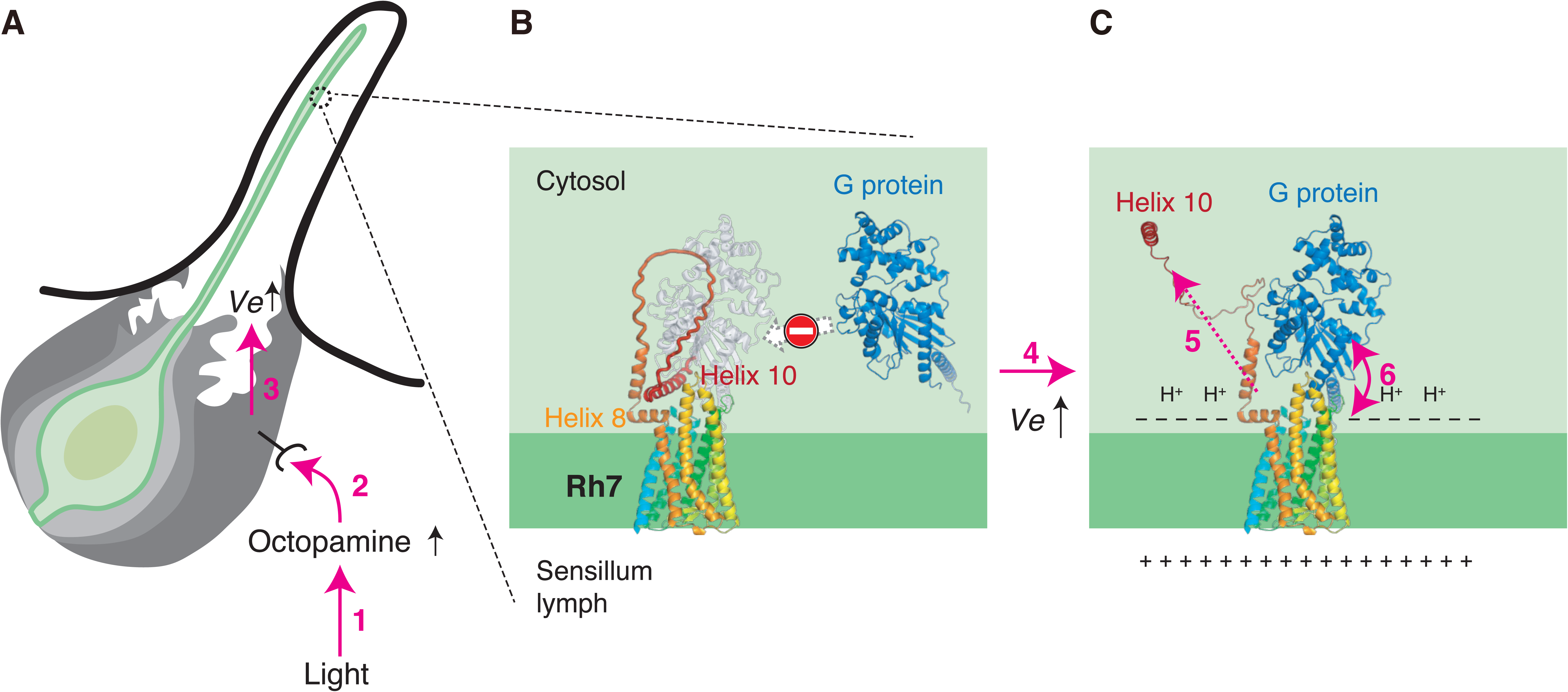
Schema of the mechanism underlying the light-evoked firing rate changes in the OSNs. Based on the structural and electrophysiological analyses presented here, the following mechanism for the light-evoked firing rate increases in the OSNs is proposed. Prior to the light stimulation, the binding of G protein to Rh7 remains inhibited by helix 10 and the long loop region of Rh7 (***A*,*B***). Light stimulation increases the concentration of octopamine in the hemolymph within the antenna (1). Octopamine then acts on auxiliary cells (2), elevating the extracellular field potential (*Ve*) within the sensillum (3). The rise in *Ve* (4) increases a potential gradient across the cell membrane, resulting in accumulation of negative charges on the intracellular surface of the cell membrane (***C***). This would trigger protonation of helix 10, resulting in release of helix 10 from helix 8 (5). Such structural changes likely allow G protein to bind to Rh7 (6) and activate second messengers to change the excitability of the OSN.

Amputation of the antennal nerve abolished the firing rate increases in the OSNs during the light stimulations (Ikeda et al., 2022). This indicates that octopamine that diffuses from outside the antenna does not evoke the firing rate increases, since octopamine in hemolymph can diffuse into the antenna even the antennal nerve is amputated. This study has not revealed whether the ephaptic transmission from the photoreceptor cells in retinae directly changes the firing rate of the octopaminergic neurons that terminate in the antenna. Busch et al. (2009) reported three types of octopaminergic neurons that innervate the antennal nerve. Whereas they do not terminate in the optic lobe and laminae, they project to the ventrolateral protocerebrum, the visual center in the protocerebrum, where the light information might be relayed to the octopaminergic neurons.

In moths, the concentration of octopamine in the brain oscillates with the circadian rhythm (Linn et al., 1996), and octopamine application changes the transepithelial potential (LFP) and enhances the pheromone response in the olfactory sensillum (Linn and Roelofs, 1986; Dolzer et al., 2001). These studies, in conjunction with our findings, indicate that octopamine, by acting on the non-neuronal auxiliary cells surrounding the OSNs, regulates the extracellular field potential to fine-tune the excitability of the OSNs to adapt to the outside environment. The theory that non-neuronal cells contribute to sensory responses as well as play an important role in homeostasis in the nervous system is also supported by a report that auxiliary cells in the antennal sensillum mediate the olfactory response to ammonia by expressing the ammonium transporter (Menuz et al., 2014). Recent study also showed that light sensation by cryptochrome expressed in the auxiliary cells and glial cells in the antenna changes the firing rates of olfactory sensory neurons (Thakur et al., 2025).

Octopamine promotes wakefulness in *Drosophila* (Crocker and Sehgal, 2008). The auxiliary cells maintain a transepithelial potential between the sensillum lymph and hemolymph by generating a high potassium concentration in the sensillum lymph with potassium pumps (Kaissling and Thorson, 1980). Hence, the light-evoked LFP increase may be mediated by a change in concentration of ions such as potassium. In mice, arousal was linked to elevations of the extracellular potassium concentration (Ding et al., 2016). It would be intriguing to examine whether non-neuronal cells commonly regulate LFP and modulate the excitability of a population of neurons under the control of biogenic amines.

*Rh7* is a unique non-visual rhodopsin gene conserved among arthropods (Senthilan and Helfrich-Förster, 2016) and functions in circadian photoentrainment in *Drosophila melanogaster* (Ni et al., 2017). While Rh7 activates Gq-type G proteins similarly to other rhodopsins in *Drosophila* (Sakai et al., 2017), Rh7-specific structures such as its longer extracellular N-terminus and intracellular C-terminal tails and relatively broad absorption spectrum compared to Rh1-6 suggested it has additional functions (Senthilan and Helfrich-Förster, 2016; Sakai et al., 2017). According to the present analysis, the C-terminal tail likely functions as a voltage-dependent gate for the α subunit of Gq-type G proteins to allow access to its binding site on Rh7. Although antennal transcriptome analyses have detected Rh7 expression (Menuz et al., 2014), Rh7 expression in the antenna was below the level detectable with anti-Rh7 antibodies (data not shown). Senthilan et al. (2019) suggested that Rh7 whose expression level is below the detection limit of the antibodies may be sufficient to induce electrophysiological responses in brain cells. The gate may allow neurons to respond to even a small change in the extracellular potential by amplifying the signal with second messengers downstream of the G protein, enabling the animals to respond to sensory stimuli in a context-dependent manner. Non-visual opsins are also expressed in the vertebrate nervous system (Yamashita et al., 2014). It would be intriguing to examine whether these non-visual opsins are also correlated with the neuronal response to the LFP changes.

## Conclusion

Rh7 is indispensable for regulating the firing rates of neurons in response to an extracellular LFP change mediated by octopaminergic neurons. Although extracellular LFP activity has generally been considered a side effect of neuronal spiking, our study reveals that the nervous system possesses an active mechanism that controls the LFP in response to the change in sensory input, resulting in alternation of behavioral patterns as well as neural firing patterns in a context-dependent manner. These findings provide valuable insights into the complex interplay between the extracellular LFP and behavioral responses, offering a foundation for further investigations into the dynamic regulation of neuronal functions.

## Data availability statement

Data and sources will be available from the corresponding author upon request.

## Author Contributions

All authors had full access to all the data in the study and take responsibility for the integrity of the data and the accuracy of the data analysis. Study concept and design: MK, KS, KI, HI, and NKT. Acquisition, analyses, and interpretation of experimental data: MK, KI, and NKT. Built the protein model: KS, HI, NKT.

## Acknowledgements

This work was supported by JST PRESTO (N.K.T), JSPS KAKENHI (JP23H04963 to K.S.; JP20H03217 and JP23H02444 to H.I.; 22770068, 24120509, 26830026, 20K06733, and 25K09696 to N.K.T.) and Interdisciplinary Computational Science Program in CCS, University of Tsukuba (K.S.). We would like to thank Fumika Hamada, Akiko Sato, Takahiro Chihara, Kazuhiko Kume, Taro Ueno, Ryusuke Niwa, Tadao Usui, Tadashi Uemura, Leslie Vosshall, Barry Dickson, Sarah Certel, Kyoto *Drosophila* Genetic Resource Center, and Bloomington *Drosophila* Stock Center for providing fly strains, and Taishi Yoshii, Craig Montell, Charlotte Förster, and Developmental Studies Hybridoma Bank at the University of Iowa for antibodies, and Aki Ejima for sharing courtship chambers. We appreciate Ayumi Tanaka and Ryoichi Tanaka for measuring light intensities and Makoto Mizunami, Tomomi Tsunematsu, Takahiro Yamashita, and Michael Schleyer for helpful comments on the manuscript.

## Supplementary Figure

**Supplementary Figure 1. Alignments of Rh7 and Rh1.**

Rh7 contains three helices (helices 8 - 10) in the C-terminal tail, whereas Rh1 has only two helices (helices 8 and 9). The transmembrane domains and DRY motif based on Senthilan and Helfrich-Förster (2016) are indicated by black and magenta lines, respectively. Glu and Arg analyzed in **Fig. 6*C*,*D*** are shown in bold.

**Supplementary Figure 2. Atomic coordinates of Rh7, Rh1, and Gq-type G protein.**

Atomic coordinates of Rh7 (A; AAF49949.2), Rh1 (B; NP_524407.1), and Gq-type G protein (C; CG17759) of *D. melanogaster*. Each column represents atom serial number (a), atom name in accordance with the Chemical Component Dictionary (b), amino acid residue name (c), chain ID (d), residue sequence number (e), x orthogonal LJ coordinate (f), y orthogonal LJ coordinate (g), z orthogonal LJ coordinate (h), occupancy (i), temperature factor (j), and element symbol (k) (from left to right).

**Supplementary Figure 3. The odor responses recorded from the ab2 sensillum.**

***A*,** The response of ab2B to hexanol (10^-3^) during sustained stimulation with ethyl acetate in wild type in a constant light condition. Left panel shows the firing rates in each bin of 300 ms. Magenta bar represents 1.2 s of hexanol puff, whereas black bar indicates the timing of 3 s puff of three different concentrations of ethyl acetate (0-10^-5^). Error bar indicates the standard error of the mean. Right graph shows the firing rates of ab2B during the period in which it showed responses to hexanol (for 1.2 s from 0.6 s after the onset of hexanol puffs). ***B*,** The response of ab2A to ethyl acetate. Firing rates of ab2A were not significantly different between wild type (WT) and *Rh7^0^* mutants during ethyl acetate stimulation, just prior to the onset of the air puff containing hexanol. ***C*,** Mean amplitudes of the LFP deflections during ethyl acetate stimulation, just prior to the onset of the air puff containing hexanol, were not significantly different between WT and *Rh7^0^* mutants. Each gray dot represents a different animal, and magenta indicates the mean ± the standard error of the mean. For statistical analyses, Kruskal-Wallis tests followed by Dunn’s multiple comparisons were performed. *, *p* < 0.05. N.S., not statistically significant.

