## Supplementary Fig 1 for "Rhodopsin 7 is indispensable for regulating the firing rates of olfactory sensory neurons in response to extracellular field potential changes in *Drosophila melanogaster*"

```
Rh7     MEAIIMTTLPNLTTDAGDSSFWLTGALSLSEMLANSSHSHSTGSTTSTAGSSATESSAVN
Rh1     -----------------------------------------------------MESFAVA
                                                               ** **

                                                                 TM1
Rh7     VGKDHDKHVNDSVSTGLSNYSNYPSYIHYRDKYDLSYIAKVNPFWLQFEPPKSSTFLIMA
Rh1     AAQLGPHFAPLSNGS-------------VVDKVTPDMAHLISPYWNQFPAMDPIWAKILT
        ..:   :..   * .:                **    .    :.*:* ** . ..    *::

                                                  TM2
Rh7     ALYCLISVVGCVGNAFVIFMFANRKSLRTPANILVMNLAICDFLMLIK-CPIAIYNNIKE
Rh1     AYMIMIGMISWCGNGVVIYIFATTKSLRTPANLLVINLAISDFGIMITNTPMMGINLYFE
        *   :*.::.  **..**::**. ********:**:****.** ::*.   *:   *   *

                  TM3                    DRY                 TM4
Rh7     GPALGDIACRLYGFVGGLSGTCAIGTLTAIALDRYNVVVHPLQPLRRCSRLRSYLIILLI
Rh1     TWVLGPMMCDIYAGLGSAFGCSSIWSMCMISLDRYQVIVKGMAG-RPMTIPLALGKIAYI
         .** : * :*. :*.  * .:* ::  *:****:*:*: :   *  :   :   *  *

                                                TM5
Rh7     WCYSFLFAVMPALDIGLSVYVPEGFLTTCSFDYLNKEMPARIFMALFFVAAYCIPLTSIV
Rh1     WFMSSIWCLAPAFGWSR--YVPEGNLTSCGIDYLERDWNPRSYLIFYSIFVYYIPLFLIC
        *  * ::.: **:. .    ***** **:*.:***:::  .* :: :: : .* ***  *

                                                   TM6
Rh7     YSYFYILKVVFTASRIQ--------------SNKDKAKTEQKLAFIVAAIIGLWFLAWSP
Rh1     YSYWFIIAAVSAHEKAMREQAKKMNVKSLRSSEDAEKSAEGKLAKVALVTITLWFMAWTP
        ***::*: .* : .:                *:. : .:* *** :. . * ***:**:*

                            TM7                         Helix 8          Helix 9
Rh7     YAIVAMMGVFGLERHITPLGSMIPALFCKTAACVDPYLYAATHPRFRVEVRMLFYGRGVL
Rh1     YLVINCMGLFKFEG-LTPLNTIWGACFAKSAACYNPIVYGISHPKYRLALKEKCP-CCVF
        * ::   **:* :*   :***.::   * *.*:*** :* :*. :**::*: ::      *:

                                                                Helix 10
Rh7     RRVSTTRSSYMTRSRSSFTHRLRTSTTGEGGMGDHRMENYLMNNNLMMVPEETEENEEIV
Rh1     GKVDDGKSSDAQSQATASEAESKA------------------------------------
         :*.  :**    . ::   . ::

Rh7     VVAEINNSISSVMEQSKF
Rh1     ------------------
```

### Supplementary Fig. 1
