## Supplementary figures and images for "Rhodopsin 7 is indispensable for regulating the firing rates of olfactory sensory neurons in response to extracellular field potential changes in *Drosophila melanogaster*"

### Supplementary Fig 3

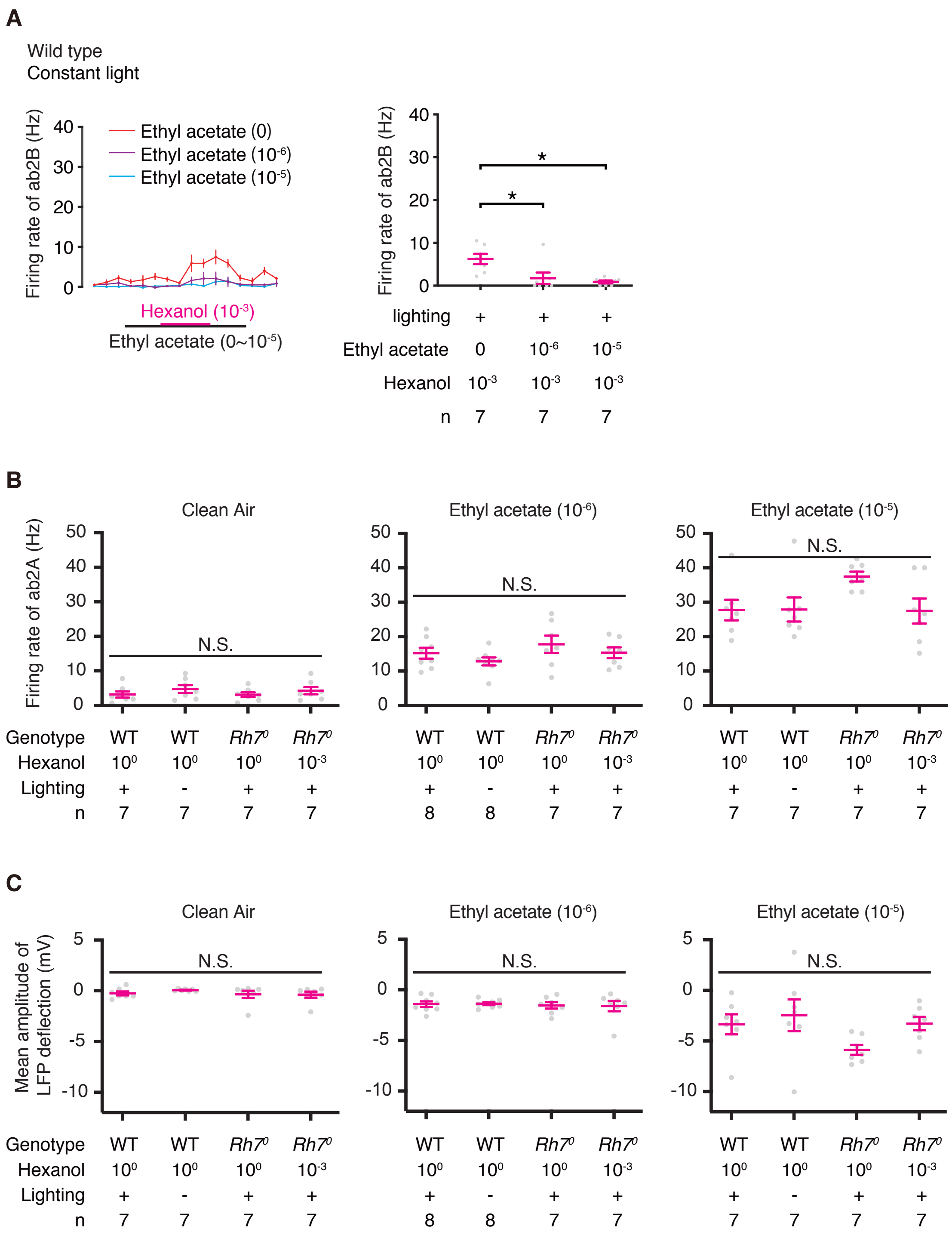


**Supplementary Fig. 3**
